# Mesoscale cortical dynamics reflect the interaction of sensory evidence and temporal expectation during perceptual decision-making

**DOI:** 10.1101/552026

**Authors:** Ivana Orsolic, Maxime Rio, Thomas D. Mrsic-Flogel, Petr Znamenskiy

## Abstract

How sensory evidence is transformed across multiple brain regions to influence behavior remains poorly understood. We trained mice in a visual change detection task designed to separate the covert antecedents of choices from activity associated with their execution. Widefield calcium imaging across dorsal cortex revealed fundamentally different dynamics of activity underlying these processes. While signals related to execution of choice were widespread, fluctuations in sensory evidence in the absence of overt motor responses triggered a confined activity cascade, beginning with transient modulation of visual cortex, followed by sustained recruitment of secondary and primary motor cortex. The activation of motor cortex by sensory evidence was selectively gated by animals’ expectation of when the stimulus was likely to change. These results identify distinct activation profiles of specific cortical areas during decision-making, and show that the recruitment of motor cortex depends on the interaction of sensory evidence and expectation.

## INTRODUCTION

As animals form judgments about the sensory scene, information represented in sensory cortical areas influences motor actions by engaging a distributed network of sensorimotor pathways. Neural correlates of decision-making have been identified across modalities and species (Hanks and Summerfield, 2017), through recordings targeting individual brain areas (Hanes and Schall, 1996; Shadlen and Newsome, 1996; Roitman and Shadlen, 2002; Romo *et al.*, 2002; Raposo, Kaufman and Churchland, 2014), or probing multiple areas serially and/or simultaneously (De Lafuente and Romo, 2005; Hernández *et al.*, 2010; Hanks *et al.*, 2015; Siegel, Buschman and Miller, 2015; Allen *et al.*, 2017; Scott *et al.*, 2017; Gilad *et al.*, 2018; Steinmetz *et al.*, 2018; Zatka-Haas *et al.*, 2019). Despite this progress, however, we still lack a mechanistic understanding of the neural processes underlying the commitment to a choice of action.

Perceptual decisions involve the interaction of sensory information with subjects’ expectations and prior knowledge leading up to behavioral choice (Gold and Shadlen, 2007; Summerfield and de Lange, 2014). Attributing neuronal responses to these pre-decision processes is challenging because they are inherently correlated with subsequent motor execution-related signals (Murakami and Mainen, 2015), which have a widespread impact on neural activity. Specifically, behavioral choice is represented across the neocortex once animals commit to a particular action (Allen *et al.*, 2017) and influences the representation of sensory stimuli (Nienborg and Cumming, 2009), while neural correlates of task-related and spontaneous overt behaviors dominate global brain activity (Musall *et al.*, 2018; Stringer *et al.*, 2018).

To separate pre-decision processes from activity related to motor execution we designed a task that allowed us to probe independently the influence of both sensory information and expectation on neural activity, while controlling the animals’ motor output. The task required mice to lick for reward in response to sustained changes in the speed of a noisy drifting grating stimulus. Mice were encouraged to respond as soon as they detected the change by restricting the window when the reward was available. Since speed changes were often ambiguous, their timing variable, and the trial difficulty randomized, mice had to continuously monitor the sensory stimulus during an extended baseline period preceding the change. During this period we observed neural activity related to pre-decision processes by measuring neural responses to stimulus speed fluctuations (Huk and Shadlen, 2005; Hanks *et al.*, 2015) free from confounds related to motor execution. Importantly, by manipulating animals’ expectation of when sustained changes in speed were likely to occur, we determined how the responses to the same stimulus speed fluctuations were influenced by temporal expectation.

Using this task we identified the behavioral strategy used by the mice to detect sustained changes in stimulus speed, showing that they combine stimulus information on the timescale of hundreds of milliseconds with their prior expectation of the timing of changes. Using wide-field calcium imaging of the dorsal neocortex we identified a cascade of activity induced by fluctuations in stimulus speed in the absence of overt motor responses. Such fluctuations triggered transient responses in visual areas and culminated in sustained activation of motor areas. The transmission of sensory information to motor cortex depended on the animals’ experience of the task and was specifically modulated by their temporal expectation of stimulus change. This localized pre-decision cascade contrasted with the widespread emergence of action-related signals associated with the execution of behavioral choice.

## RESULTS

### Visual change detection task

We trained head-fixed, food-restricted mice in a visual change detection task, which required them to lick for reward in response to a sustained increase (from here referred to as *change*) in the speed of a grating stimulus (Figure 1A). Mice were placed on a styrofoam cylinder and were required to remain stationary for a random interval of 3 to 8.8 seconds to initiate a trial. At the onset of each trial, mice were presented with the baseline grating stimulus drifting upward or downward, with a mean temporal frequency (TF) of 1 Hz. During this baseline period, movements of the cylinder in either direction (exceeding 2.5 mm in a 50 ms window) aborted the trial. The temporal frequency of the grating stimulus varied around the mean every 50 ms (3 monitor frames) during both the baseline and the change periods on 70% of trials (referred to as *noisy* trials, Figure 1B). Such noisy trials provided a window to determine the strategies that mice might use to perform the task as well as to probe stimulus evoked modulation of cortical activity during decision making (Huk and Shadlen, 2005; Hanks *et al.*, 2015). After a randomly chosen delay period the mean TF increased, and the mice were required to lick within a response window of 2000 ms to receive a drop of soy milk reward. The frequency of correct licks (‘hits’) depended on the magnitude of the TF change, with mice reliably detecting increases in TF to 2 Hz (>91 % detected) or 4 Hz (>95 % detected; Figure 1C, 19734 noisy trials, 109 sessions, 6 mice). Reaction times were also modulated by the magnitude of stimulus changes, with mice responding more swiftly to larger increases in TF (Figure 1D-E).

**Figure 1.**
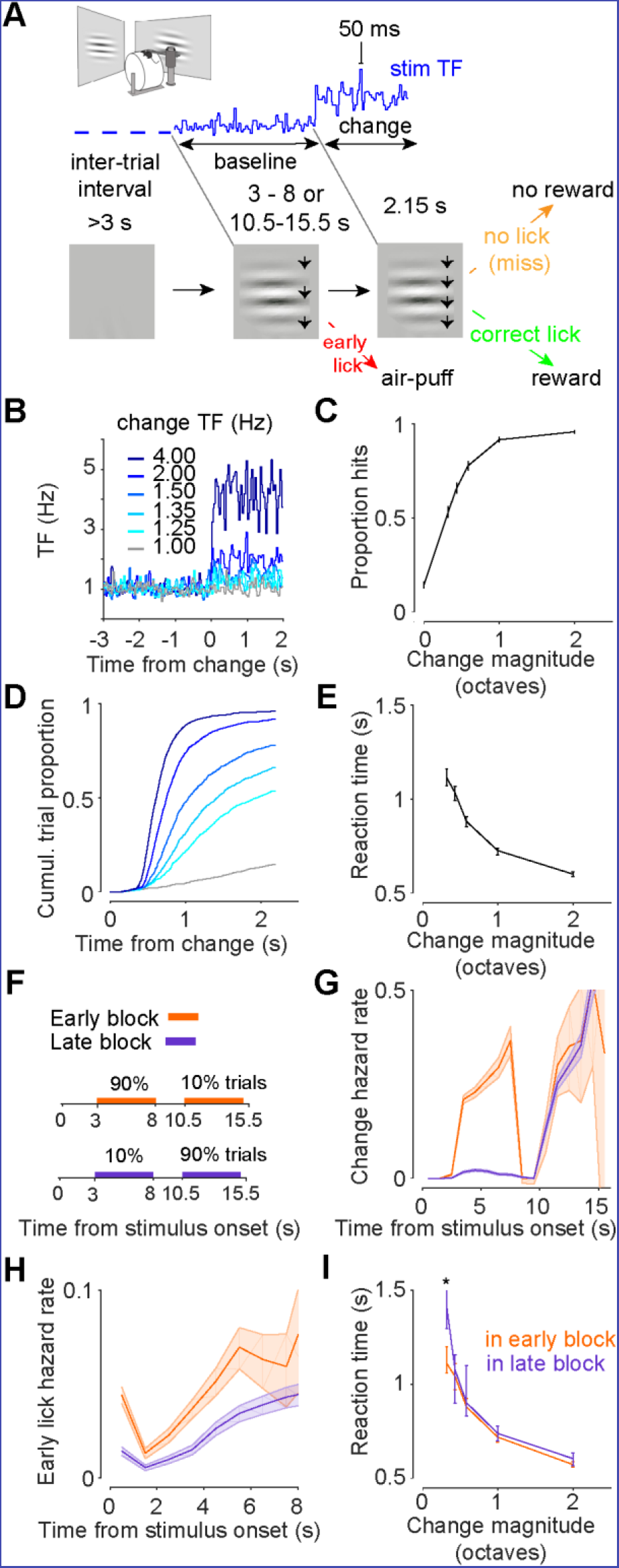
Change detection behavioral task and animal performance. **A**. Experimental setup and change detection task structure. Mice were head-fixed, and two monitors were placed on each side of the animal. Square patch of a drifting sine grating appeared after a randomized inter-trial interval. After randomized time, TF of the stimulus increased. If the mouse detected the change (correct lick – green) reward of soy milk drop was given. If the mouse licked before the change, an air-puff was delivered (early lick – red). **B**. Example stimuli aligned to change onset. Darker color corresponds to larger change magnitudes. **C**. Detection rate is modulated by change magnitude (6 mice, error bars are 95% CI). **D**. Cumulative distribution of reaction times across stimulus changes. Colors as in B. **E**. Median reaction times are modulated by change magnitude (6 mice, error bars are 95% CI). **F**. Timing of changes across early and late change blocks. G. Probability of changes as function of time (hazard rate) in early and late blocks. H. Early lick hazard rate is modulated by anticipation of change. **I.** At low change magnitudes, responses to early changes (3-8 s after stimulus onset) are slower in late blocks, when changes are not expected (* – p < 0.01, Wilcoxon rank sum test).

On ~18% of trials (3590/19734) mice licked prior to change during the baseline period, which aborted the trial and was penalized with an air puff to the cheek. To explore whether the timing of such early licks was influenced by animals’ expectation of when changes might occur, we varied the distribution of change times during the trial in blocks (Figure 1F-G). In early change blocks, changes occurred between 3 and 8 s after the stimulus onset on most trials (90 %) and between 10.5 and 15.5 s on probe trials (10 %). The timings were reversed in late change blocks. The timing of early licks depended on the hazard rate of stimulus changes that the mice experienced (probability of changes given that no change has occurred and the trial has not been aborted, Figure 1G). Early lick hazard rate was elevated at the start of the trial in early blocks (Figure 1H), when changes were more frequent. Mice were also faster to respond to the most difficult change during this period if the change was expected (Figure 1I). Therefore, prior expectation of when changes might occur influenced animals’ decisions to lick.

### GP classification model uncovers stimulus features driving mouse behavior

Several behavioral strategies could explain the features of mouse behavior described above. For example, mice might decide to lick by integrating visual signals, or by detecting fast outliers in the noisily drifting stimulus. Distinguishing between these strategies based on trial-average statistics is challenging (Brunton, Botvinick and Brody, 2013). To understand how sensory evidence is transformed into a decision to lick, we took advantage of the stochastic fluctuations in TF on noisy trials and examined the TF content of baseline stimuli preceding early licks by computing the lick-triggered average stimulus. Early licks were preceded by increases in temporal frequency spanning the period of ~250-1000 ms prior to lick onset (Figure 2A). This observation shows that sensory information over this epoch contributes to animals’ behavior but does not unambiguously reveal how subjects weigh evidence in their decisions to lick (Okazawa *et al.*, 2018).

**Figure 2.**
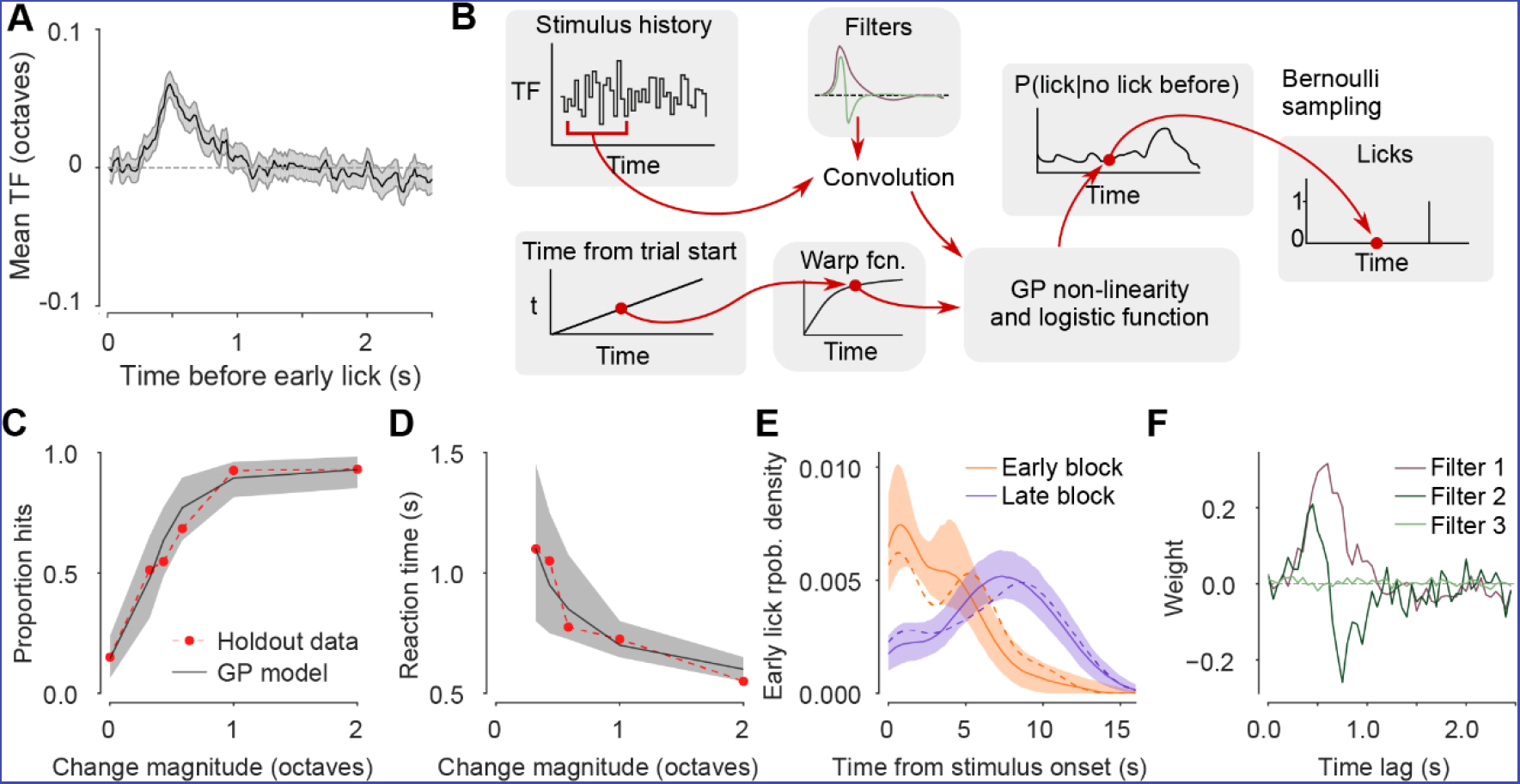
Gaussian process classification model reveals animals’ behavioral strategy. **A**. Average stimulus preceding licks during the baseline stimulus (n=6 mice, shading is 95% CI). B.Structure of the Gaussian process classification model. **C-E.** Model performance for an example mouse (425 holdout trials). The model captures the animal’s detection performance (**C**) and reaction times (**D**), as well as the timing of early licks (**E**, computed using a Gaussian kernel density estimate; holdout data – dashed lines; model predictions – solid line and shading). Early and late block trials are combined in panels **C** and **D**. To fit and evaluate the model, 14944 behavioral trials were assigned to training, validation, and test subsets (8966, 2989, and 2989 trials respectively). Of these 1277, 424, and 425 trials were recorded during hazard rate manipulation sessions in stationary mice. During the remaining trials, mice were free to run during stimulus presentation. F. Principal filters learned during model optimization. The first two filters reveal the main stimulus features sufficient to capture mouse behavior.

To answer this question, we developed a statistical model optimized to predict the momentary lick hazard rate during sustained changes in stimulus speed as well as during baseline periods, based on the history of visual input over the past 2.5 s and time elapsed since the start of the trial (Figure 2B). To identify features of the stimulus that drove animals’ choices, we restricted stimulus information available to the model to a low-dimensional linear projection of stimulus history defined as the convolution of stimulus history with a set of filter vectors. Filter outputs and elapsed time were combined by a Gaussian process (GP) non-linearity to estimate the log-odds of licking during each sample of the task. The model assumes that the contributions of time and stimulus information to the log-odds of licking are additive, which is equivalent to combining current sensory evidence with prior odds of licking based on timing. Although this model has no direct biological interpretation, it provides an unbiased description of how mice transform stimulus information and time since trial start into licks, akin to linear-nonlinear-Poisson models used to characterize neuronal receptive fields. The model accurately captured the trial-average statistics of mouse behavior, including psychometric and chronometric curves (Figure 2C-D), as well as the timing of early licks during the trial (Figure 2E). The full model outperformed models that received stimulus or timing inputs alone (Figure S1).

We next examined the stimulus filters, whose weights and number were optimized during model training (Figure 2F). The shapes of the top two filters were consistent across all mice (N=6) whose behavior we quantified (Figure S2). The weights of both filters were close to zero for time lags of 0 to 250 ms, a period equivalent to non-decision time, reflecting sensory and motor delays unrelated to decision-making. The first filter, whose larger amplitude indicates a greater role in predicting mouse behavior, resembled the lick-triggered average stimulus and had large positive weights at time lags of ~250 to ~1000 ms and small negative weights at lags of 1000-2000 ms (Figure 2F, S2). It is therefore sensitive to sustained increases in the TF of the grating over baseline. The second filter was almost symmetric and resembled a derivative filter, with positive weights between ~250 and ~500 ms and negative weights between ~500 and ~1000 ms (Figure 2F, S2D,H,L,P,T). It is therefore sensitive to abrupt changes in TF of the grating. The weights of the remaining filters were close to zero with the exception of the third filter in 2 of 6 mice, which resembled the derivative filter but was shifted in time (Figure S2L,P).

These analyses show that mouse behavior is best explained as a combination of both integration of TF on the timescale of ~1 second and outlier detection strategies, with greater weight given to the former, as well as by the expectation of when the sustained changes in stimulus speed might occur.

### Imaging activity dynamics across dorsal cortex during the task

We next systematically characterized the patterns of neural activity underlying the processing of sensory signals and their transformation into putative motor commands across dorsal cortex. To accomplish this, we imaged transgenic mice expressing GCaMP6s (Wekselblatt *et al.*, 2016) (11130 trials, 47 sessions, 6 mice) using a low magnification epifluorescence microscope that allowed us to simultaneously capture calcium signals across the entire dorsal surface of the mouse neocortex (Figure 3A). To compensate for changes in fluorescence arising from hemodynamic changes, we interleaved illumination at 470 nm and 405 nm, close to the isosbestic point of GCaMP6s, and used frames acquired at 405 nm to correct calcium traces for fluorescence fluctuations unrelated to neural activity (Allen *et al.*, 2017). Imaging was conducted at 50 Hz, resulting in a hemodynamics-corrected frame rate of 25 Hz. In parallel, we monitored animals’ pupil diameter, as well as facial and body movements. To identify the imaged brain areas at the end of each imaging experiment, we reconstructed whole brain volumes using serial two-photon tomography and registered them to the Allen Common Coordinate Framework (CCF v3, Figure S3). We then used superficial blood vessel patterns to align functional imaging data to the serial tomography volumes and define cortical area boundaries based on the Allen CCF annotation (Figure 3A, Figure S4).

**Figure 3.**
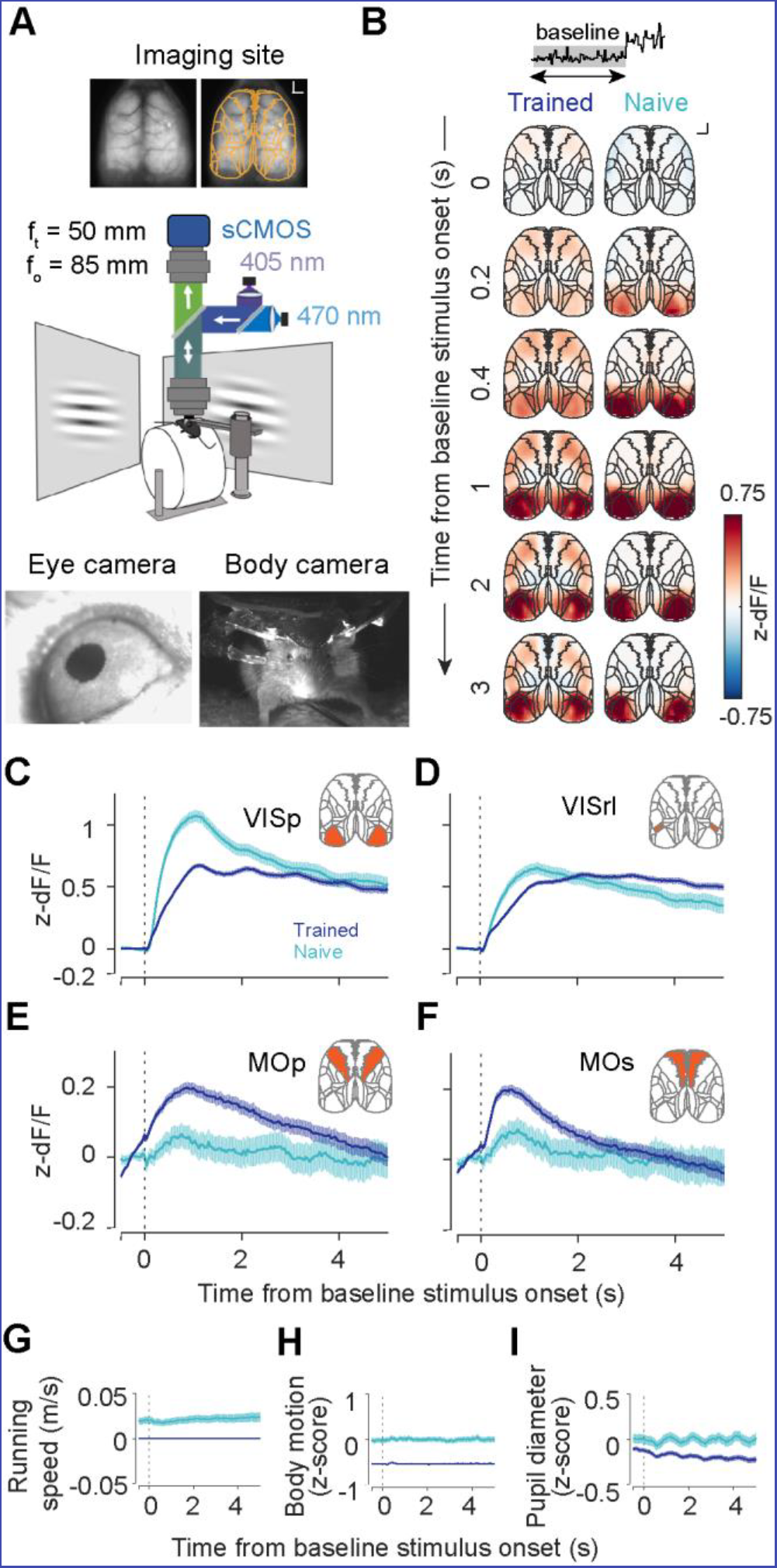
Stimulus onset broadly activates dorsal cortex in trained animals. **A**. Widefield imaging experimental setup. Top – extent of the imaging site; orange – borders of regions of interest (ROI) analyzed after brain registration. Outer borders were cropped to the extent of the imaging site. Centre – schematic of the behavioral setup and wide-field macroscope. Bottom left – example image from eye camera. Bottom right - example image from camera placed in front of the animal to capture facial and body movements. **B**. Average z-scored fluorescence response during baseline stimulus period in trained (left, 6 mice) and naïve (right, 3 mice). Inset – shading indicates the analyzed trial epoch. Scale bar – 1 mm. **C-F.** Mean z-scored responses of selected ROIs for trained and naïve animals. Vertical line marks the stimulus onset. Inset – shaded region corresponds to ROI shown: VISp – primary visual cortex (**C**), VISrl – rostrolateral visual area (**D**), MOp – primary motor cortex (**E**), MOs – secondary motor cortex (**F**). **G-I.** Mean traces of running speed (**G**), body motion (**H**), pupil diameter (**I**) for trained and naive animals.

In the presentation of our results, we focus on responses in four cortical areas, which show markedly different patterns of activity during baseline and change periods – primary and rostrolateral visual areas VISp and VISrl, and primary and secondary motor areas MOp and MOs. Responses in all imaged ROIs are presented in supplemental figures and movies.

### Visual stimulus onset engages a distributed cortical network in trained mice

We first analyzed the patterns of activity evoked by the onset of the baseline stimulus (Figure 3B). In mice performing the task, grating onset triggered sustained activation of primary and secondary visual areas (Figure 3C-D, 6631 noisy trials longer than 1.5 seconds, 47 sessions, 6 mice; Supplemental Movie 1) followed by recruitment of secondary and primary motor cortex (Figure 3E-F). While grating onset triggered responses of similar or even larger magnitude in visual areas in naïve mice, responses in motor areas were markedly weaker (1680 noisy trials from 10 sessions in 3 mice; Supplemental Movie 1). Thus, recruitment of motor cortex by the onset of the visual stimulus occurs even in the absence of movement (Figure 3G-I) and depends on animals’ experience of the task.

### Action-related signals are represented throughout dorsal cortex

We next examined the patterns of activity evoked by sustained changes in TF of the grating that the mice were trained to detect. Change onset triggered an increase in fluorescence across the dorsal surface of the cortex on hit trials (Figure 4A, D, 1974 noisy trials; Supplemental Movie 2). These responses occurred earliest in motor areas, with latency of 480 ms for the strongest stimuli (secondary motor area – MOs; time to half-max response) compared to 600 ms latency for primary visual cortex – VISp (Figure S5A). On more difficult trials response latencies followed the increase in reaction times. Across stimulus strengths, the time course of neural responses followed the movement of the mouse, as captured by the body camera (Figure 4D, S5B-C). This widespread modulation of cortical activity was not observed on miss trials (Figure 4H, S5D-E, 463 noisy trials) suggesting that these responses are related to the execution of animals’ choices in the task.

**Figure 4.**
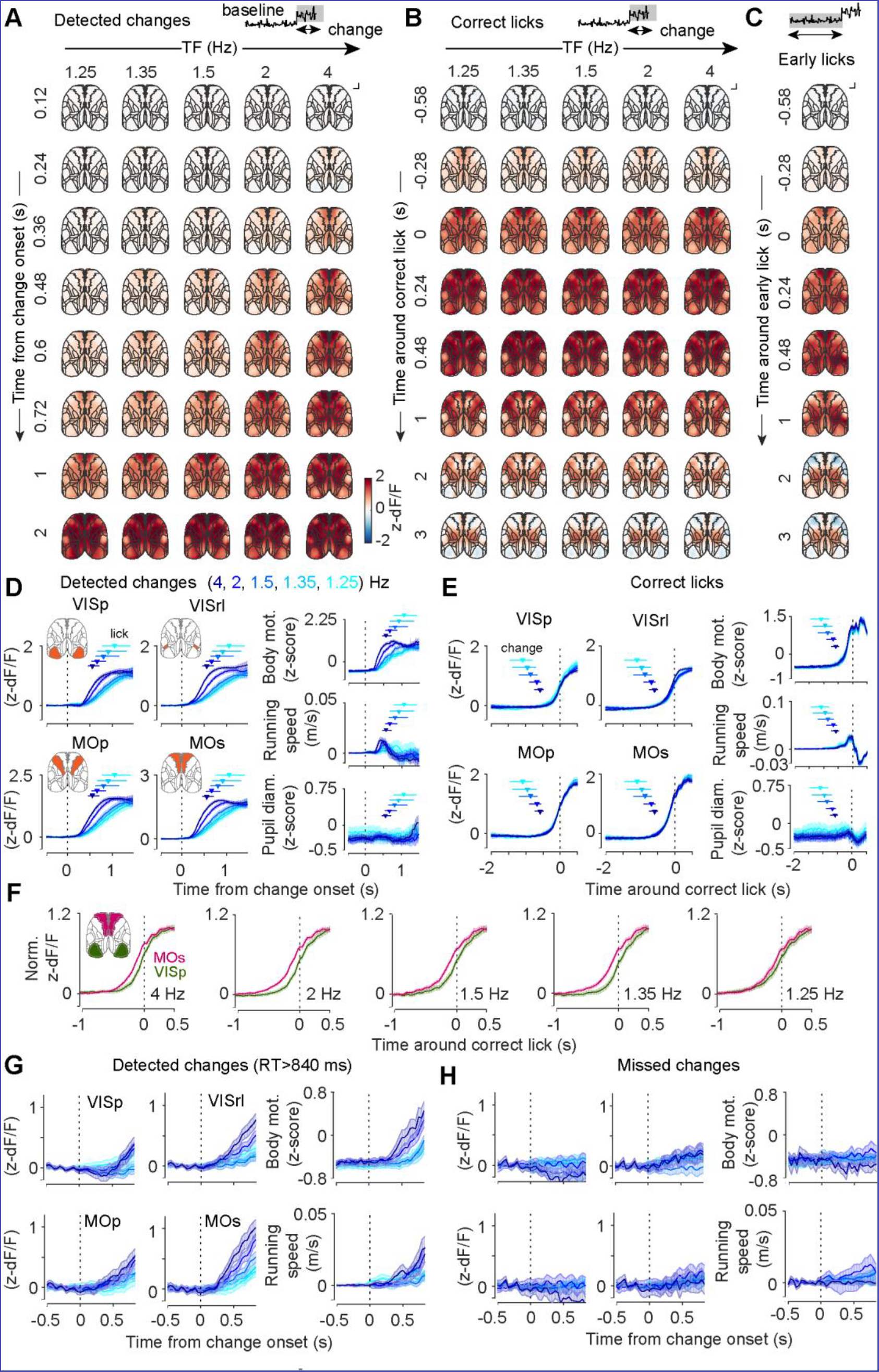
Responses across dorsal cortex during the change period are global and dominated by licking-related activity. **A-C.** Maps of mean z-scored fluorescence across the dorsal cortex aligned to change onsets on hit trials (**A**), correct licks (‘hits’) (**B**) and early licks (**C**). Panels **A** and **B** are sorted by change strength. Inset – shaded is the trial epoch from which data is represented. **D-E.** Mean z-scored fluorescence of selected ROIs and behavioral measures across change strengths aligned to change onsets on hit trials (**D**) and aligned to correct lick onsets (**E)**. Shading is 95% CI. Horizontal lines and markers represent the interquartile range and the median reaction times (**D**) or stimulus change times (**F**). **F.** Pre-lick activity of secondary motor cortex precedes activity in primary visual cortex (normalized to peak activity for each ROI). **G-H.** Mean z-scored fluorescence of selected ROIs and behavioral measures across change strengths aligned to change onsets on hit trials where lick happened at least 840 ms from the change point (**G**) and miss trials (**H**). Body movements preceding licks accompany increase in activity on hit trials.

When aligned to the onset of licking, fluorescence responses were stereotyped across stimulus strengths (Figure 4B,E, S6A-C, Supplemental Movie 3) and were similar to responses aligned to licks that occurred before the change (Figure 4C, S6D-E, 1564 noisy trials). While lick-related activity was global, it did not appear synchronously across the cortex. It was detectable earliest in secondary motor cortex, anterior visual and midline areas (VISal – 360 ms prior to lick; VISam, VISa, VISrl, MOs, RSPd, ACAd – 320 ms; quantified as the time to cross 10% of maximum response on 1.5 Hz change trials) followed by primary motor cortex, somatosensory cortex, and primary visual cortex (SSptr, RSPv – 280 ms; MOp, SSpul, SSpll, VISpm – 240 ms; VISp, SSpn – 200 ms; SSpm – 160 ms; Figure 4F and Supplemental Movie 3).

Lick-aligned activity dominated but did not fully account for cortical responses following the onset of sustained changes in stimulus speed. To illustrate this, we examined activity on hit trials with long reaction times (>840 ms). During these trials, change onset triggered a gradual increase in fluorescence, which was modulated by the strength of the stimulus (Figure 4G). However, while no licks were present during this period, this activity was correlated with other overt movements preceding licking as captured by the body camera (Figure 4G) and was absent on miss trials when no licks or other movements occurred (Figure 4H). Together, these analyses show that the activation of the dorsal cortex following the onset of stimulus speed changes reflects a combination of responses related to licking and other body movements, which may mask activity associated with the covert processes underlying the decision to lick.

### Fluctuations in sensory evidence trigger a localized cascade of activity from visual to motor areas in absence of overt movements

The extended baseline period, during which mice had to monitor the visual stimulus without overt changes in their behavior, made it possible to examine the patterns of neural activity associated with transformation of sensory information without the confounds related to execution of movement. To characterize the temporal progression of visual stimulus processing in the dorsal cortex, we quantified the effect of sensory evidence during the baseline stimulus on widefield fluorescence at different time lags using linear regression

(Figure 5A-B, S6, 1039391 frames from 6894 trials, Supplemental Movie 4). To ensure that movement-related activity immediately preceding licks did not affect this analysis, we excluded fluorescence frames from trials interrupted due to early licks or movement acquired less than 1 second prior to these events, as well as frames following change onset.

**Figure 5.**
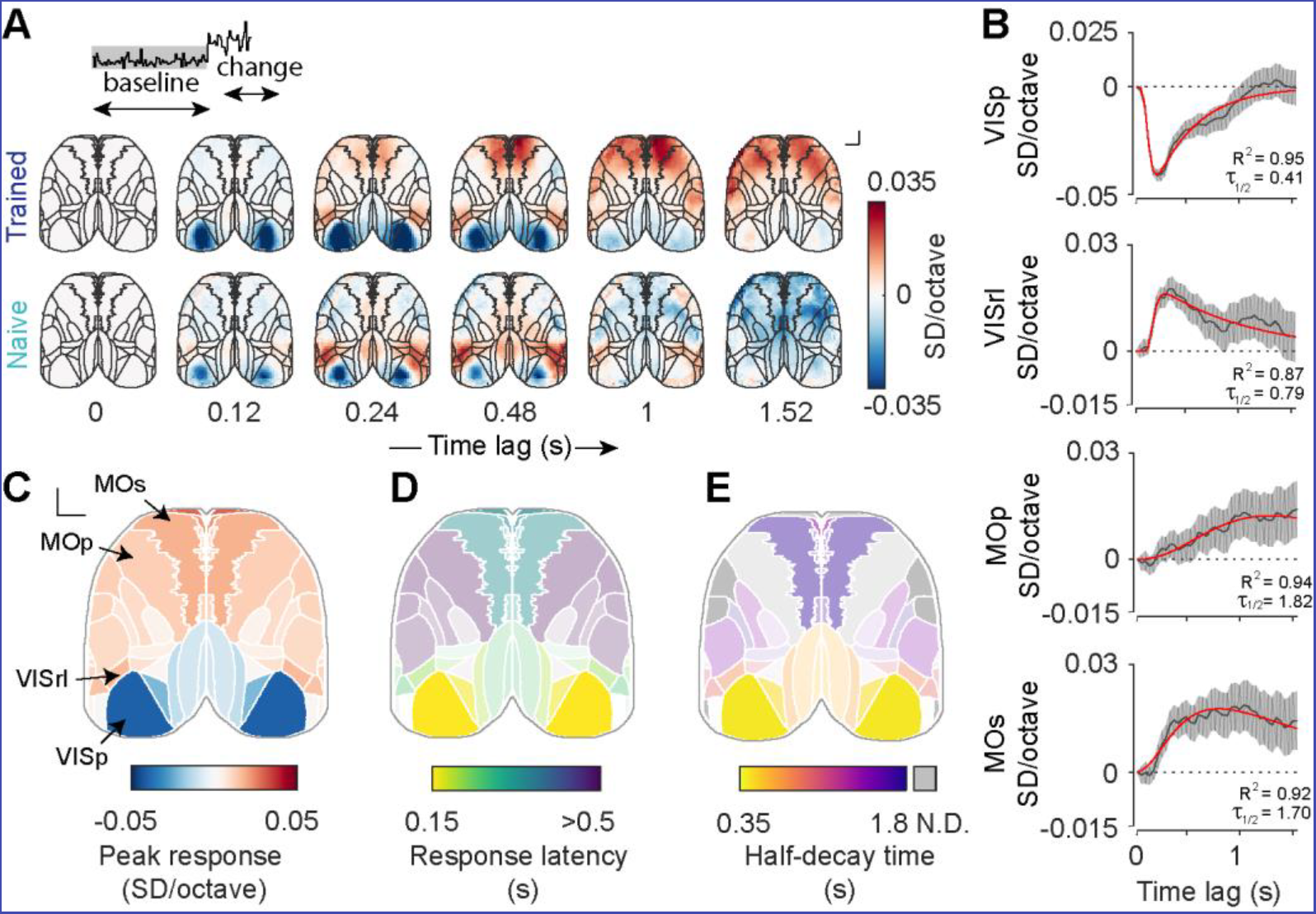
Subthreshold stimulus fluctuations trigger a localized cascade of activity across dorsal cortex. **A**. Maps of regression coefficients of widefield fluorescence against baseline stimulus TF in trained (top) and naïve (bottom) animals across time lags. **B**. Time course of regression coefficients of widefield fluorescence against baseline stimulus TF in example ROIs; regression coefficients (gray, 95% CI) and multiexponential fit (red). **C**. Peak response across recorded brain regions. D. Response latency (time to half maximum) across brain regions. Saturation is scaled based on response magnitude (panel **C**). **E**. Response half-decay time across brain regions. Saturation is scaled based on response magnitude (panel **C**). Regions for which half-decay time could not be determined are gray.

Temporal frequency of the baseline stimulus was negatively correlated with activity of primary visual cortex but positively correlated with activity of anterior higher visual areas (Figure 5A-C). The latencies (defined as time until 50% of maximum response; Figure 5D) were shortest in primary visual cortex (VISp – 147 ms) and higher posteromedial visual area (VISpm – 159 ms), followed by anterior higher visual areas: rostrolateral visual area (VISrl – 209 ms) including adjacent regions of somatosensory cortex, anterior visual area (VISa – 217 ms) and anterolateral visual area (VISal – 250 ms). Since widefield imaging captures local population activity (Makino *et al.*, 2017), this modulation is consistent with the typical preference of primary visual cortex and higher visual area neurons. While VISp neurons tend to prefer slow visual speeds, neurons in areas VISal, VISrl, and VISa preferentially respond to high speeds (Andermann *et al.*, 2011; Marshel *et al.*, 2011). Similar responses in visual areas were also present in naive mice, consistent with their sensory-driven origins (Figure 5A).

In trained mice, the modulation of visual areas by temporal frequency was followed by activation of secondary motor cortex (MOs - 333 ms) and weaker recruitment of primary motor cortex (MOp – 632 ms) and somatosensory areas (lower and upper limb SSp-ll, SSp-ul, 561-904 ms; nose and mouth SSp-n, SSp-m, ~686-869 ms). Unlike responses in visual cortical areas, the modulation of motor cortical activity was not observed in naive mice (Figure 5A, 362291 frames from 1680 trials, Supplemental Movie 4).

Cortical areas also differed in the offset dynamics of their responses. Activity in primary visual cortex rapidly decayed to baseline (half-decay time of 410 ms, Figure 5B,E), suggesting that it is largely modulated by the immediate history of sensory stimulation. In contrast, responses in higher visual areas and motor areas were sustained (VISrl – 790 ms, VISal – 930 ms, and VISa – 970 ms) with half-decay times exceeding 1.5 s in secondary and primary motor cortices (MOs – 1700 ms, MOp – 1820 ms, Figure 5B,E, Figure S6). The step-like impulse response of secondary motor cortex suggests that its activity reflects integration of sensory evidence.

However, given the nature of widefield imaging, we cannot discern if these sustained responses are carried by one homogeneous population or if they are mediated by different populations with different time courses of activity (Scott *et al.*, 2017).

Measurements of offset dynamics are limited by the temporal resolution of calcium signals. An estimate of this resolution is provided by responses in primary visual cortex, which decay with a half-life of 410 ms (Figure 5B). This value is similar to the reported half-decay time for somatic signals in GCaMP6s transgenics (Dana *et al.*, 2014).

The regression analysis described above revealed the sign and time course of modulation of the activity of dorsal cortex by the visual stimulus. To characterize this relationship in more detail, we computed mean responses to the extremes of the stimulus during the baseline period, which carry different information for the animal – fast (‘pro-licking’, N = 41194) and slow (‘anti-licking’, N = 42253) stimulus pulses (1.5 standard deviations above or below the mean TF, respectively) using responses to stimulus near the mean TF (+/-0.5 standard deviations, N = 467681 pulses) as a reference (Figure 6A; Supplemental Movie 5). In primary visual cortex and area VISpm, fast stimulus pulses were associated with a decrease in fluorescence compared to the reference stimulus response, while slow stimulus pulses triggered an increase in fluorescence (Figure 6B, Figure S4). These effects were reversed in anterior higher visual areas VISrl, VISal, and VISa (Figure 6B, Figure S4). In contrast, motor areas responded selectively to fast stimulus pulses, while slow pulses had no significant effect (Figure 6B). Thus, secondary and primary motor cortex were activated specifically by stimulus fluctuations that mice were trained to detect.

**Figure 6.**
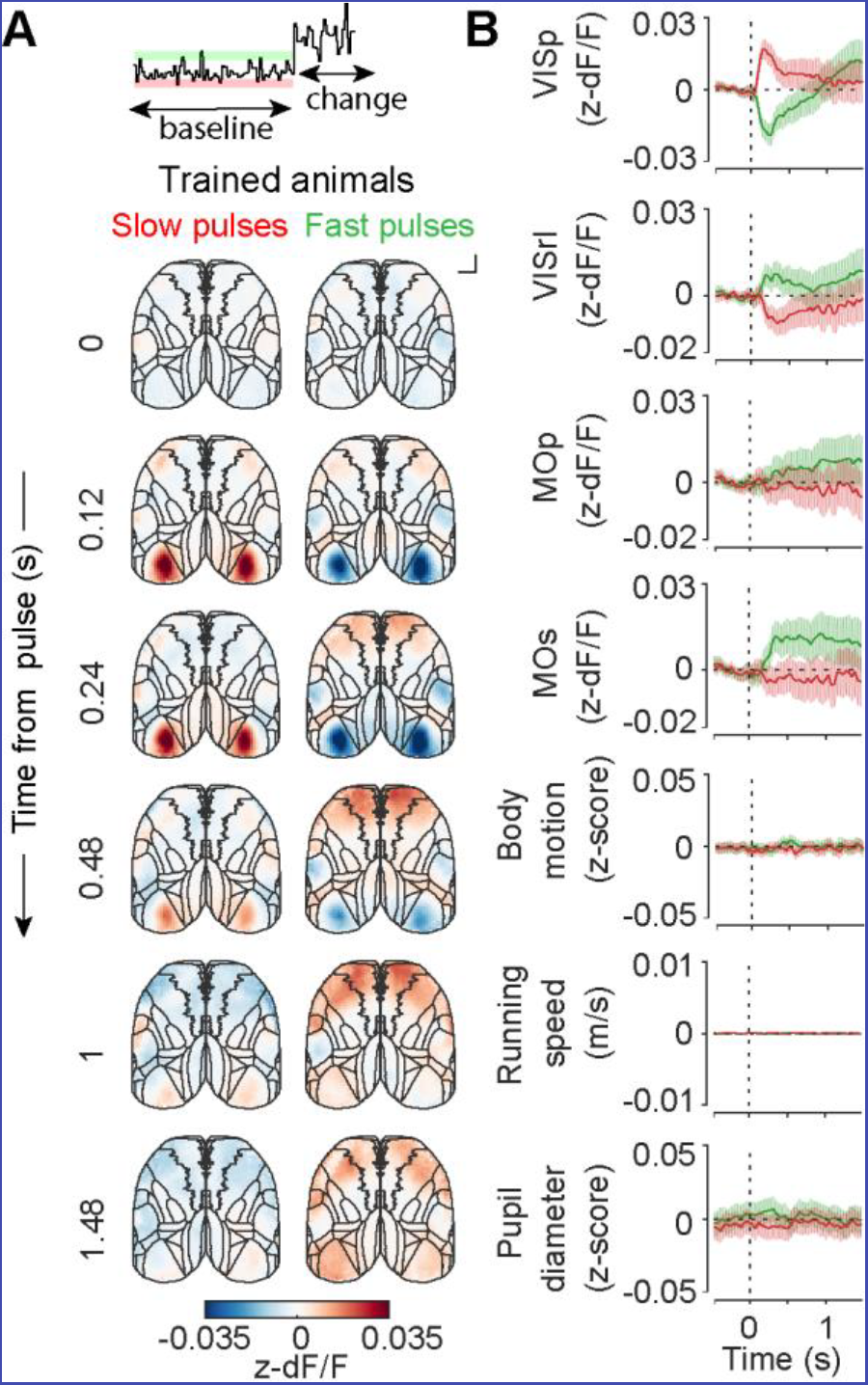
Responses to anti- and pro-licking subthreshold stimulus fluctuations across dorsal cortex. **A**. Maps of mean z-scored fluorescence responses to anti-(slow) and pro-licking (fast) subthreshold stimulus fluctuations in trained mice (6 mice). **B**. Mean z-scored fluorescence of selected ROIs and behavioral measures aligned to anti-(slow, red) and pro-licking (fast, green) subthreshold stimulus fluctuations in trained mice. Shading is 95% CI.

In the analyses described above we took care to minimize the impact of movement-related activity on our estimates of neural responses. Importantly, neither fast nor slow baseline stimulus pulses produced consistent overt motor responses, as measured by the body camera (Figure 6B). In a different version of the task, mice were required to run on the wheel to initiate a trial but were free to modulate their running speed during stimulus presentation (performance of mice during running sessions in Figure S8, 82005 trials, 281 sessions, 6 mice). Although this was not explicitly encouraged by the task, we found that mice changed their running speed in response to the baseline grating stimulus, speeding up after slow stimulus pulses and slowing down after fast stimulus pulses (Figure S9C-D). The resulting correlation between baseline stimulus TF and running speed confounded the interpretation of widefield fluorescence responses. In contrast to the localized cascade of activity we observed in the stationary version of the task, baseline stimulus fluctuations in running mice were followed by global modulation of dorsal cortical activity (Figure S9A-B, S10, Supplemental Movie 6). The time course of these global responses resembled that of running behavior but was opposite in sign (Figure S10). These observations highlight the importance of controlling for movement in the interpretation of neural data and motivated us to focus our analyses on the stationary version of the task.

### Temporal expectation gates the activation of motor cortex by sensory evidence

When making decisions, animals take advantage of immediate sensory evidence as well as their predictions of the environment, but how the cortex combines these signals is unknown. To answer this question, we determined how the processing of sensory evidence in the dorsal cortex was influenced by animals’ temporal expectation of stimulus change. We compared the relationship between stimulus speed and widefield fluorescence in the periods where the animal was expecting a change to occur (early change blocks) and those when the change was not expected (late change blocks) using linear regression as above (Figure 7A; 244491 imaging frames from 2521 trials in early block and 521827 imaging frames 4373 trials in late block). We focused on the period when the early lick hazard rate differed between blocks (first 6 seconds after the baseline stimulus onset, Figure 1H).

**Figure 7.**
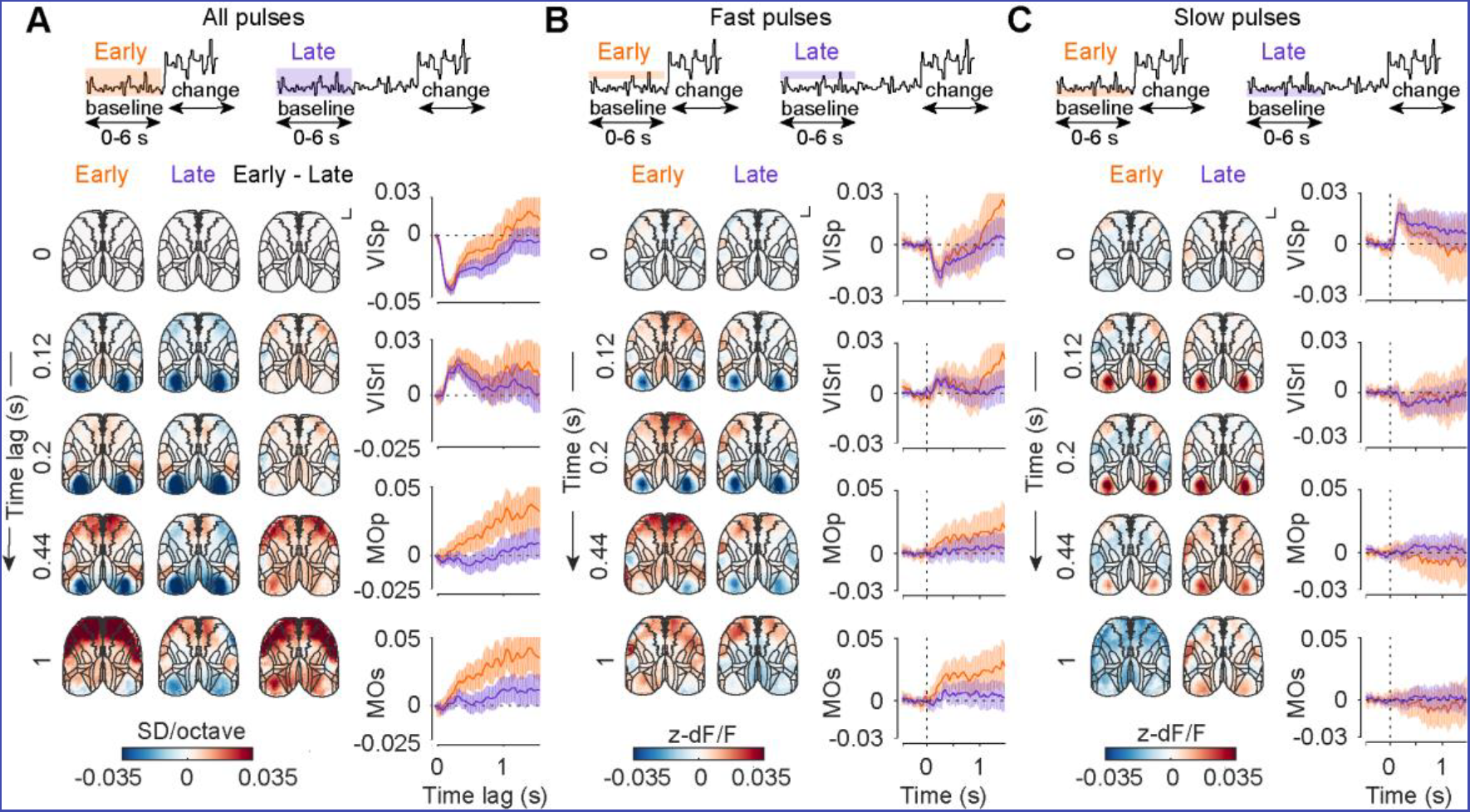
Temporal expectation modulates responses to stimulus fluctuations in motor but not sensory areas. **A.** Regression coefficients of widefield fluorescence against baseline stimulus temporal frequency during 0-6 s of the trial in early (orange) and late (purple) change blocks, and difference between the blocks (early – late). Maps (left) and selected ROIs (right). Shading is 95% CI. **B-C.** Mean z-scored fluorescence responses to pro-(fast, **B**) and anti-licking (slow, **C**) subthreshold stimulus fluctuations during 0-6 s of the trial in early and late change blocks. Notation as in **A**.

We found that animals’ temporal expectation of stimulus speed change specifically modulated the relationship between stimulus speed and fluorescence in motor areas (Figure 7A). Both secondary and primary motor cortices responded more strongly to stimulus fluctuations during early blocks, wherein changes were expected soon after the start of the trial. In contrast, the initial responses to sensory evidence in visual areas were indistinguishable between early and late blocks.

To understand how animals’ temporal expectation affected the processing of sensory evidence in motor cortex, we computed mean responses to fast (pro-licking) and slow (anti-licking) stimulus pulses during the first 6 s of the trial in early and late change blocks (Figure 7B; Early change block: fast pulses: N = 9655; slow pulses N = 9957; reference pulses: N = 110282. Late change block: fast pulses: N = 20864; slow pulses: N = 21111; reference pulses: N = 234372). Responses to slow stimulus pulses were similar between the two change blocks. On the other hand, responses to fast stimulus pulses in motor areas were enhanced when animals were expecting the change. Thus, temporal expectation selectively enabled the transfer of task-relevant sensory information to secondary and primary motor cortex.

## DISCUSSION

### Visual change detection as a paradigm to study perceptual decisions

In order to study the neural correlates of variables leading up to perceptual decisions, we developed a visual change detection task wherein mice had to observe a drifting stimulus whose speed fluctuated noisily to detect sustained changes in speed. We identified the strategy used by mice in the task using a combination of model-based and model-free approaches. First, by examining the stimuli preceding licks during the baseline stimulus, we showed that stimulus information over ~1 second contributed to animals’ decisions to lick. Second, we used a Gaussian process classification model to identify stimulus features that could predict animals’ choices on a moment-by-moment basis. This analysis suggested that mouse behavior is best explained as a combination of integration and outlier detection strategies. Finally, by manipulating the timing of changes during the trial, we showed that animals’ expectation of when stimulus speed changes might occur contributed to their decision to lick.

Typically, neural activity in reaction time tasks reflects the interaction of multiple concurrent and correlated signals, including those related to sensory integration, action selection and execution (Park *et al.*, 2014). The baseline period of our task allowed us to independently characterize the patterns of neural activity underlying the processing of sensory evidence separate from the responses associated with the execution of behavioral choice (Figure 8). By taking advantage of the stochastic nature of stimulus speed during the prolonged baseline period, we uncovered the patterns of cortical activity preceding the commitment to a decision, reflecting the transformation of sensory evidence and its interaction with animals’ expectation in the absence of overt motor responses.

**Figure 8.**
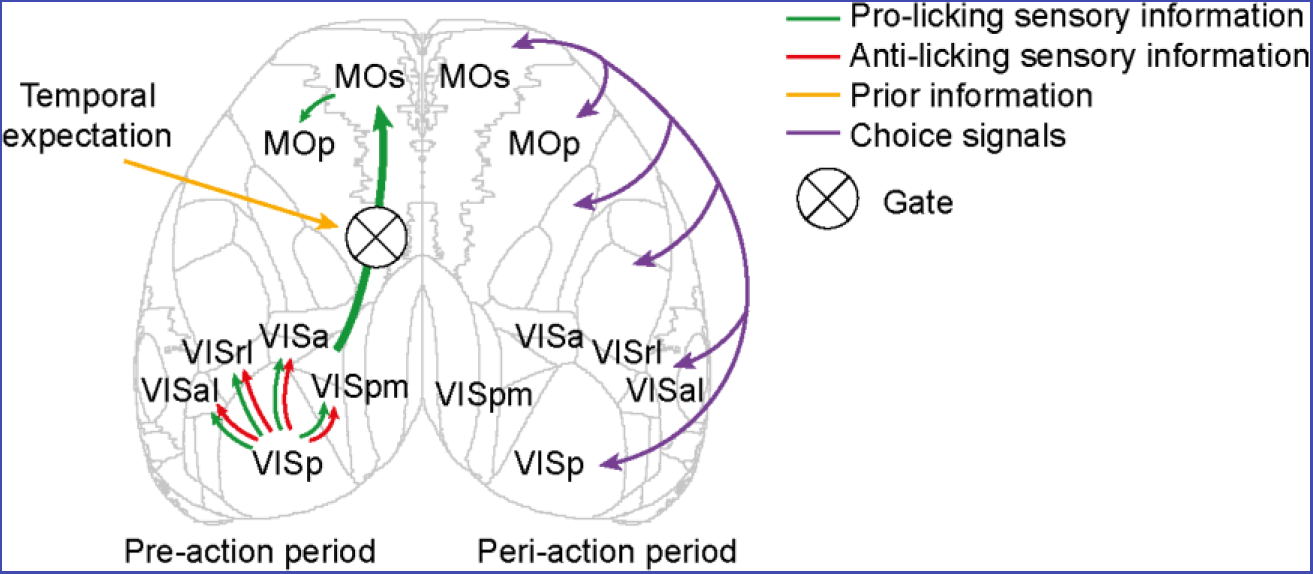
Proposed functional flow of sensory and choice-related signals in mouse dorsal cortex. Sensory information modulates activity in primary and higher visual areas. Only pro-licking sensory signals are transmitted to secondary motor cortex, where they persist on prolonged timescales. This functional flow from visual sensory areas to secondary motor cortex is gated by animals’ expectation. When the animal executes its choice, movement-related preparation signals are broadcast globally across the dorsal cortex.

### Localized cascade of cortical activity reflects pre-decision processing of sensory evidence in the absence of overt motor responses

We found that fluctuations in stimulus speed during the baseline period specifically modulated several areas of dorsal cortex. This modulation differed in sign and temporal dynamics across areas. Tracking the ongoing stimulus, visual cortical areas VISp and VISpm responded *transiently* and bidirectionally to pro-licking (high speed) and anti-licking (low speed) stimulus information with short latencies. In anterior visual areas VISa, VISal, and VISrl, which form the core of the mouse posterior parietal cortex (Hovde *et al.*, 2018), bidirectional responses to fluctuations in the visual stimulus speed were *sustained* over hundreds of milliseconds. This observation is consistent with electrophysiological recordings in rats, which suggest that posterior parietal cortex faithfully represents accumulated sensory evidence (Hanks *et al.*, 2015). Activity of visual cortical areas contrasted with that of secondary motor cortex, which responded selectively to pro-licking information in a sustained manner, with signals persisting longer than 1 second. While sensory responses in primary and secondary visual cortical areas were present in both trained and naïve mice, the transfer of sensory information to motor cortex occurred only in trained animals. Thus, acquisition of the task shaped the flow of sensory information in the dorsal cortex.

The localized nature of pro-licking selective responses in secondary and primary motor cortex is consistent with systematic perturbations of activity across the cortex in a tactile discrimination task, which identified an area of anterior motor cortex as uniquely required for preparation and execution licking responses (Guo *et al.*, 2014). Furthermore, the rat homologue of the secondary motor cortex contains the frontal orienting field (FOF), which has been proposed as a key locus in the evolution of orienting decisions (Erlich, Bialek and Brody, 2011). In a task in which rats base their choices by integrating auditory signals over hundreds of milliseconds (Brunton, Botvinick and Brody, 2013), FOF represents evolving behavioral choices (Hanks *et al.*, 2015) and is required for task performance (Erlich *et al.*, 2015). Inactivation studies indicate that MOs plays an important role in the expression of behavioral choices in perceptual tasks (Guo *et al.*, 2014; Allen *et al.*, 2017; Zatka-Haas *et al.*, 2019). Sustained modulation of MOs activity by sensory input in our task reveals that it is engaged even in the absence of overt motor responses and thus has a role beyond motor execution. Together, these observations suggest that this area contributes to perceptual choices independent of sensory modality or motor readout, perhaps by representing an emerging plan of action.

### Temporal expectation gates recruitment of motor cortex by sensory evidence

Our behavioral analyses revealed that temporal expectation of *when* the stimulus speed might change influenced animals’ behavior in the task. Widefield imaging permitted us to identify the stage of the sensory processing cascade, where temporal expectation had an impact on neural responses. Contrary to earlier studies in rodents (Jaramillo and Zador, 2011), we did not observe temporal expectation-related modulation in sensory areas. On the other hand, secondary and primary motor areas preferentially responded to pro-licking stimulus fluctuations in periods when the stimulus speed change was likely. Thus, temporal expectation influenced the transmission of sensory evidence from visual to motor areas. While emergence of such selective transmission with learning has been previously implied (Makino *et al.*, 2017), our results suggest that it can also be modulated dynamically in the absence of overt movements and provide the behavioral framework to further study the neural bases of this modulation across multiple brain regions.

A key question is how task-relevant inputs in visual cortex are relayed to motor areas. Our experiments cannot disambiguate whether these signals are transmitted through direct corticocortical or indirect subcortical pathways. Projections from sensory cortex to the basal ganglia have been implicated in mediating the influence of sensory evidence on perceptual decisions (Znamenskiy and Zador, 2013). A cortico-basal ganglia loop may play a similar role in our task, by relaying task-relevant visual signals to motor areas in an expectation-dependent manner.

### Widespread movement-related modulation of dorsal neocortex

We found that the behavioral responses in the task had a global influence on neural activity. Even prior to the onset of licking, movements recorded by the body camera were accompanied by widespread recruitment of dorsal cortex (Figure 4G), potentially masking localized neural signals underlying the decision to lick. Such global influence of motor behavior on cortical activity is consistent with recent reports (Allen *et al.*, 2017; Musall *et al.*, 2018; Stringer *et al.*, 2018). However, the extent to which these global signals arise as the result of preparation or execution of movements, or of sensory feedback associated with them, remains unclear. If these signals indeed represent a form of efference copy, and broadcast the selected motor action throughout the cortex, they may serve as a substrate for reinforcement learning (Fee, 2014).

The ubiquity and magnitude of movement-related signals poses a major challenge for interpretation of neurophysiological signals, when motor behavior is poorly controlled. This challenge is illustrated by the version of the change detection task, in which mice were free to run on the wheel during presentation of the baseline stimulus. Mice were more likely to stop after increases in the speed of the stimulus, even though this aspect of their behavior was not explicitly reinforced in the task. The resulting correlation between sensory evidence and running speed masked the effects of the stimulus on cortical activity, confounding them with running speed related modulation (Figure S9-10).

The extended baseline period and stimulus design in our task provide a way to capture neural activity resulting from the interaction of sensory signals and animals’ expectation free of global modulations associated with changes in overt behavior such as running or licking. The stark differences between the patterns of neural activity underlying the processing of sensory signals and those arising during the execution of licks highlight the importance of task designs that disambiguate these related and often concurrent processes.

## ACKNOWLEDGEMENTS

We thank Ruben Portugues, Santiago Jaramillo, and Athena Akrami for comments on earlier versions of the manuscript, Robert A. A. Campbell for serial two-photon tomography protocols and code for brain registration, Lisa Hoermann for help with behavioral training and histology, Fabia Imhof for help with histology, Devon Cowan for development of custom parts for head-fixation, Adil G. Khan for contributing code for RT state machine implementation, SWC high-performance computing facility for computational resources, Raymond Strittmatter and Biozentrum mechanical workshop for custom opto-mechanical parts, Robert Häring and Biozentrum electronic workshop for custom electronic circuits. This work was supported by the Biozentrum University of Basel Fellowship for Excellence (I.O), Biozentrum Core Funds, Gatsby Charitable Foundation (GAT3212 / GAT3361) and Wellcome (090843/E/09/Z).

## AUTHOR CONTRIBUTION

I.O. and P.Z. designed the experiments; I.O. carried out the experiments; M.R. and P.Z. developed the GP classification model; I.O. and P.Z. analyzed the data; I.O., P.Z. and T.M.-F. wrote the manuscript.

## EXPERIMENTAL MODEL AND SUBJECT DETAILS

### Mice

All experiments were conducted in accordance with institutional animal welfare guidelines licensed by Swiss cantonal veterinary office. To express calcium indicator in excitatory cells throughout the cortex, we crossed heterozygous Camk2a-tTA (JAX#007004) and homozygous tetO-GCaMP6s (JAX#024742) mice (Wekselblatt *et al.*, 2016).

### Surgery

Two weeks before the start of behavioral training, mice were switched to reversed light-cycle. Environment enrichment was provided in form of a running wheel and cardboard tunnels. After acclimatization, eleven adult male mice (84 – 104 days) underwent surgery to implant a head-plate and expose the skull over the dorsal cortex for transcranial imaging. Animals were anaesthetized with a mixture of fentanyl (0.05 mg per kg), midazolam (5.0 mg per kg), and medetomidine (0.5 mg per kg). The animal's skull was exposed and cleaned, and a metal head-plate was secured to the scull around the edge of the occipital plate and the superior temporal line using dental cement (Super-Bond C&B, Sun Medical). The exposed imaging site was covered with transparent dental cement (Polymer L-Type Clear, Sun Medical) and a glass coverslip (150 um thickness), pre-cut using a diamond scribe to match the exposed surface of the skull (Silasi *et al.*, 2016). A custom-made 3D printed light shield was then cemented to the preparation.

### Behavioral setup

Behavioral setups, similar as described in (Poort *et al.*, 2015), were placed in sound isolated boxes. The mouse was head-fixed and placed on Styrofoam wheel (d = 20 cm, w = 12 cm). Wheel movements were monitored using a rotary encoder (pulse rate 1000, Kübler) coupled to the wheel axle. Two 21.5” monitors were placed on each side of the animal (~20 cm away from the animal, slightly angled and tilted towards animals’ body), covering approximately 100x70 degrees of visual space. Monitors were gamma-corrected with maximum luminance of ~40 cd/m^2^ (Konica Minolta, LS-100 Luminance Meter). Custom written software in MATLAB controlled stimulation using PsychToolbox-3 (Kleiner, Brainard and Pelli, 2007). Soy milk reward were delivered through the spout in front of the animal. Reward delivery was regulated via pinch valve (NResearch). The spout was coupled to a piezo element whose output was used to measure animals’ licking. Custom electronic hardware was used to amplify the piezo signals and control the valve. An air tube was placed ~3 cm from the animals’ right cheek to deliver light air-puffs (200 ms, 2 bar pressure, tip was cut open to 2 mm). Animals’ right eye was imaged with a CMOS camera (Imaging Source, 30 Hz) in order to tract eye movements and pupil diameter. A second camera was placed in front of the animal capturing animals’ body movements. To increase throughput of animals, animals were trained in parallel on 5 different setups. Animals were assigned to the setups randomly from sessions to session. Behavioral data were acquired using custom written code in LabView (National Instruments) and PCI-6320 acquisition card (National Instruments).

### Behavioral task

Each trial began with a grey isoluminant screen. After a randomized delay (minimum 3 s + sample from an exponential distribution with the mean 0.5 s) baseline stimulus appeared (sinusoidal grating with the spatial frequency of 0.04 cycles per degree, square patch of ~75 degrees). The temporal frequency of the baseline stimulus increased after a randomized baseline period. Change times were sampled from an exponential distribution with a mean of 4 s truncated at 5 s and added to an offset of 3 s in early blocks and 10.5 s in late blocks. Initially, the offset for early probes (early changes that occur in late block) was 4 seconds (29/109 sessions), in rest of the sessions it matched offset of the early block distribution. Removing trials outside of the 4-8 seconds overlap window during these sessions did not affect our conclusions in Figure 1I. On noisy trials, temporal frequency of the grating was drawn every 50 ms (3 monitor frames) from a lognormal distribution, such that log_2_-transformed TF had the mean of 0 and standard deviation of 0.25 octaves. The geometric mean TF on noisy trials was 1 Hz. In a subset of trials (30%) no noise was added and baseline stimulus had a constant TF of 1 Hz. Mice were trained to report increase in mean temporal frequency by licking the spout to trigger reward delivery (drop of soy milk). If mice did not lick within 2.15 s from the change, the trial was a miss trial. If mice licked before the change happened, they recieved an air puff to the cheek. Responses in the first 150 ms (‘refractory licks’, 58/19734 trials) were not rewarded and were excluded from analysis of hit trials.

### Behavioral training

Before animals underwent training on the temporal frequency change detection task, several pre-training steps were taken in order to habituate the animal to the setup.

One week after the surgery, mice were food-restricted and behavioral training started. Animals were handled for a minimum of 3 sessions, until mice were comfortable with the experimenter and were climbing on experimenters’ hand while being given drops of soy milk. Animals were then introduced to short manual restraint periods in a soft cloth after which animal was given soy milk reward. Next, animals were head-fixed and placed on the running wheel of the behavioral training setup (10 – 20 min) with the monitors turned off and were trained to run on the wheel for reward. This step typically took 4 sessions. Two mice were not trained further than this step and were assigned to the naïve cohort.

Next, to ensure that the animals understood the relationship between the stimulus presented on the monitor and reward availability, mice were pretrained on a simple task, where they had to lick in response to a change in the orientation of the grating. At this stage, translation of the grating was linked to the running speed of the mouse. As soon as mice started responding to the change in grating orientation, this step was complete. One mouse, which failed to learn to respond to orientation changes, was not trained further and was added to the naïve cohort. Eight mice proceeded training on the temporal frequency change detection task. Two of these mice were excluded from study due to lack of progress (too high abort rate due to early licks). It took the remaining six mice 14-21 sessions to learn the task. Mice were initially allowed to run during the task. After observing strong modulation of cortical activity associated with running, mice were required to be stationary during the task.

### Widefield calcium imaging

Widefield calcium imaging was carried out using a custom-built tandem-lens epifluorescence macroscope using two photographic lenses (85mm f/1.8D objective, 50mm f/1.4D tube lens, Nikon) placed in face-to-face orientation (Ratzlaff and Grinvald, 1991). Excitation light from two LEDs: 470 nm (M470L3, Thorlabs, with excitation filter FF02-447/60-25, Semrock), and 405 nm (M405L3, Thorlabs, with excitation filter FF01-405/10-25, Semrock) was combined using a dichroic (FF458-Di02-25x36, Semrock) and delivered in Koehler configuration through a dichroic mirror (FF495-Di03, Semrock) placed in the infinity focused imaging path. Average power ~0.05 mW/mm^2^, similar to that in other studies (Wekselblatt *et al.*, 2016). Images were acquired after emission filter (525/50-25, Semrock) using an sCMOS camera (pco.edge 5.5, PCO) at 50 Hz in rolling shutter mode and binned on the fly 2x2 using manufacturer software. With our lens combination this resulted in a resolution of ~20 µm per pixel. Excitation wavelengths were temporally interleaved by a microcontroller (Teensy 3.2) triggered by the camera rolling shutter exposure output. To avoid rolling shutter artefacts and crosstalk between 470 nm and 405 nm excitation frames, illumination was restricted to periods when all the lines being acquired corresponded to the same imaging frame (t_global_ in manufacturers’ documentation). A photodiode (PDA100A-EC, Thorlabs) recorded the onset of each visual stimulus frame to ensure precise alignment between visual stimulation and imaging data.

### Serial two-photon tomography

At the end of imaging experiments, mice were anaesthetized with 5 mg/kg midazolam, medetomidine 0.5 mg/kg, and 0.05 mg/kg fentanyl and five DiI (Invitrogen D3911) tracks were made across the imaging site. Tracks were made by coating the micropipette with DiI dissolved in ethanol. Locations of the DiI tracks were recorded under the widefield macroscope to ensure that imaging frames could be successfully registered to *ex vivo* brain volumes. However, since we found that blood vessel patterns could be reliably reconstructed from *ex vivo* data, DiI tracks were not used for *ex vivo* / *in vivo* registration.

The mice were then anaesthetized with sodium pentobarbital and transcardially perfused with 4% paraformaldehyde. The brains were extracted, post-fixed overnight in 4% paraformaldehyde, and stored in 50 mM phosphate buffer. Brains were coronally sectioned (100 µm steps) and imaged at two optical planes per physical section resulting in voxel size of (x = 1.32 µm y = 1.32 µm z = 50 µm) using a custom serial two-photon tomography microscope. After illumination correction and stitching, brain volumes were registered to the Common Coordinate Framework provided by the Allen Institute for Brain Science (CCF, v.3 © 2015 Allen Institute for Brain Science, Allen Brain Atlas API, available from http://brain-map.org/api/index.html) using Elastix (Klein *et al.*, 2010) by applying rigid affine transformation followed by non-rigid deformation as previously described (Han *et al.*, 2018).

To reconstruct the superficial blood vessel pattern from serial two-photon tomography volumes, we first identified the dorsal surface of the volume as the locations of the first voxel crossing a manually selected brightness threshold. We smoothed the location values with a median filter and used the fluorescence of the voxels at near this surface to reveal blood vessels (Figure S3).

### Behavioral data analysis

We included all the sessions after mice crossed the threshold of detecting more than 80% easiest changes in no-noise trials and interrupted less than 55% no-noise trials due to early licking. Average detection rate across sessions for easiest change was 96.9 ± 9.3% (mean ± sd), average early lick rate 19.4 ± 16.3% (mean ± sd). We excluded 6/115 sessions due to high abort rate due to movement. In the remaining sessions the average abort rate due to movement was 43.4 ± 17.8% (mean ± sd) where 52% of all movement induced aborts happened during the first 3.5 seconds of the stimulus.

When computing behavioral performance, all error bars are 95% confidence intervals, unless otherwise stated. For psychometric curves and hazard rates, confidence intervals were estimated using *binofit* in MATLAB, for chronometric curves they were calculated as the 0.025 and 0.975 quantiles of 2000 bootstrap samples with replacement).

To estimate hazard rates, the number of early licks and changes in one second bins was normalized by the total number of trials, excluding trials where early lick or change have already happened, or trial was aborted due to movement prior to the start of the bin.

To compute lick triggered averages, stimuli preceding early licks were averaged across animals, revealing mean stimulus information content prior to the lick. Confidence intervals were estimated by resampling early licks (2000 bootstrap samples with replacement).

### GP classification model

A Gaussian process classification model was trained to predict the lick hazard rate in discrete time samples corresponding individual TF fluctuations (50 ms). The formulation and implementation of the model are described in detail in Supplemental Information. In brief, the model received as its inputs the history of the visual stimulus over the past 50 samples (2.5 s) and time elapsed since the start of the trial. The stimulus history was filtered by multiplying the stimulus vector with a filter matrix, whose columns define the stimulus features that best predict mouse behavior. The effective dimensionality of the filtered stimulus space was controlled by placing a hierarchical Gaussian prior on each column of the filter matrix, shrinking superfluous projections to 0. The time input was passed through a non-linear monotonic warping function, to account for non-stationary nature of timing behavior. Filtered stimulus history and warped time served as inputs to the GP component of the model, whose output predicted the log-odds of licking. The covariance of the GP prior was defined as the sum of Matérn 5/2 kernels on filtered stimulus and warped time. To jointly fit behavior in different hazard rate blocks, the model was extended to include a hierarchy across experimental condition by extending the kernel to include population and hazard block-specific components. The model was implemented on the basis of the Stochastic Variational GP model class of the *gpflow* package in Python (de G. Matthews *et al.*, 2017). Computer code for model optimization and analysis of model fits can be found at https://github.com/BaselLaserMouse/rt_model_orsolic.

## Neural data analysis

### Pre-processing of imaging data and haemodynamic correction

Saved frames were checked for dropped frames and XY motion artefacts, binned 4x4 using a box kernel, and separated to 405 nm and 470 nm channels. The camera offset (average of 10000 dark frames) was removed from each frame. Each pixel in each session was low-cut filtered (cut-off at 0.00333 Hz), preserving the DC offset. Haemodynamic signal correction: Channel 405 nm was linearly interpolated to timepoints of 470 nm channel frames by taking the average 405 nm frames immediately before and after each 470 nm frame. Ratio of 470 and 405 channels was normalized by the mean of the ratio.

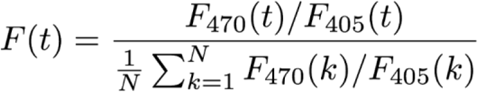

Frames were further down-sampled 2x resulting in 170 um per pixel resolution. To correct for differences in illumination and prep quality across the imaging site within individual sessions, and across sessions and animals, fluorescence traces for each pixel were normalized by their standard deviation within each imaging session. Finally, normalized imaging frames were aligned to a common reference in the Allen CCF (Figure S3).

### Responses to task events

To compute fluorescence responses associated with the baseline stimulus onset, stimulus change, correct, and early licks, we first identified the imaging frame, which was being exposed when a given event occurred. This frame corresponds to time 0 in fluorescence traces in Figures 3-4. We then extracted fluorescence traces around each event. For stimulus onset traces in Figure 3, we excluded frames acquired after change onset or less 1 second prior to early licks or wheel movement and resulting trials shorter than 0.5 s. Aligned traces were then baseline corrected by subtracting the mean fluorescence in 480 ms (for stimulus and change onset) or 2000 ms (for licks) prior to event onset.

### Responses to subthreshold TF fluctuations

We first resampled the TF of the baseline stimulus at the sampling rate of imaging acquisition. To do this, we computed the geometric mean TF presented during each imaging frame acquired during the baseline stimulus, weighted by their presentation duration:

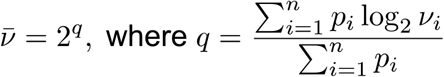

*v*_*i*_ is the TF of the i-th monitor frame presented during a given imaging frame, *p*_*i*_ is its duration (16.7 ms or less, for monitor frames, which spanned two imaging frames) and is the number of monitor frames overlapping the imaging frame.

For the regression analysis in Figure 5, we then generated a matrix of fluorescence responses to individual resampled TF fluctuations by subtracting the baseline fluorescence at the onset of the monitor frame. We then computed regression coefficients of baseline corrected fluorescence against log_2_-transformed TF for each time lag, only including fluorescence frames acquired during the baseline stimulus and at least 1 second prior to early licks or wheel movements.

We then quantified the time course of regression coefficients in different cortical areas by fitting a multiexponential model:

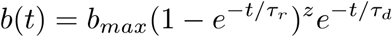

Peak response *b*_*max*_, power coefficient *z*, and rise and decay time constants τ_r_ and τ_d_ were optimized using *lsqnonlin* in MATLAB. The peak response in Figure 5C was directly given by the corresponding fit parameter. Response latency (Figure 5D) was estimated as the time lag, at which the multiexponential fit exceeded 50% of its maximum absolute value. The half decay time (Figure 5E) was estimated the as time following the response maximum, at which the fit, extrapolated if necessary, fell below 50% of its maximum value. If this did not occur within the 4 s window from stimulus pulse, half decay time is reported as not determined (N.D.).

For the analyses of responses to binned TF fluctuations in Figures 6-7, we computed mean fluorescence traces aligned to resampled TF fluctuations within each TF bin, again only including fluorescence frames acquired during the baseline stimulus and at least 1 second prior to early licks or wheel movements. To account for the overall time course of the baseline stimulus response (Figure 3), we then subtracted the mean response to the middle bin from responses to extreme bins. Due to the large sample size (tens to hundreds of thousands of imaging frames), confidence intervals were computed using the Normal approximation from the standard errors of mean fluorescence responses in each bin.

### Videography data extraction

Right eye was illuminated with a custom made IR-light source and imaged using a CMOS camera (DMK22BUC03, Imaging Source, ~30 Hz). Frames were filtered using 2-D gaussian filter (σ=2) and thresholded to low IR light reflectance areas (< 7.5 % image max intensity). Regions were filtered based on circularity (perimeter squared to area ratio < 1.6 x 4π) and size (>100 pixels). Edges of the area were detected using canny method and filtered using a Gaussian filter (σ=1). An ellipse was fitted iteratively to the region matching the criteria by minimizing the geometric distance between the area outline and the ellipse using nonlinear least squares (MATLAB function *fitellipse*, Richard Brown). Pupil diameter was estimated as the major axis of the ellipse after z-scoring within each session to correct for differences in illumination.

A second CMOS camera was placed in front of the animal capturing animals’ face and body. Body motion was expressed as the mean squared difference between the two consecutive frames z-scored within each session.

---

**Figure S1.**
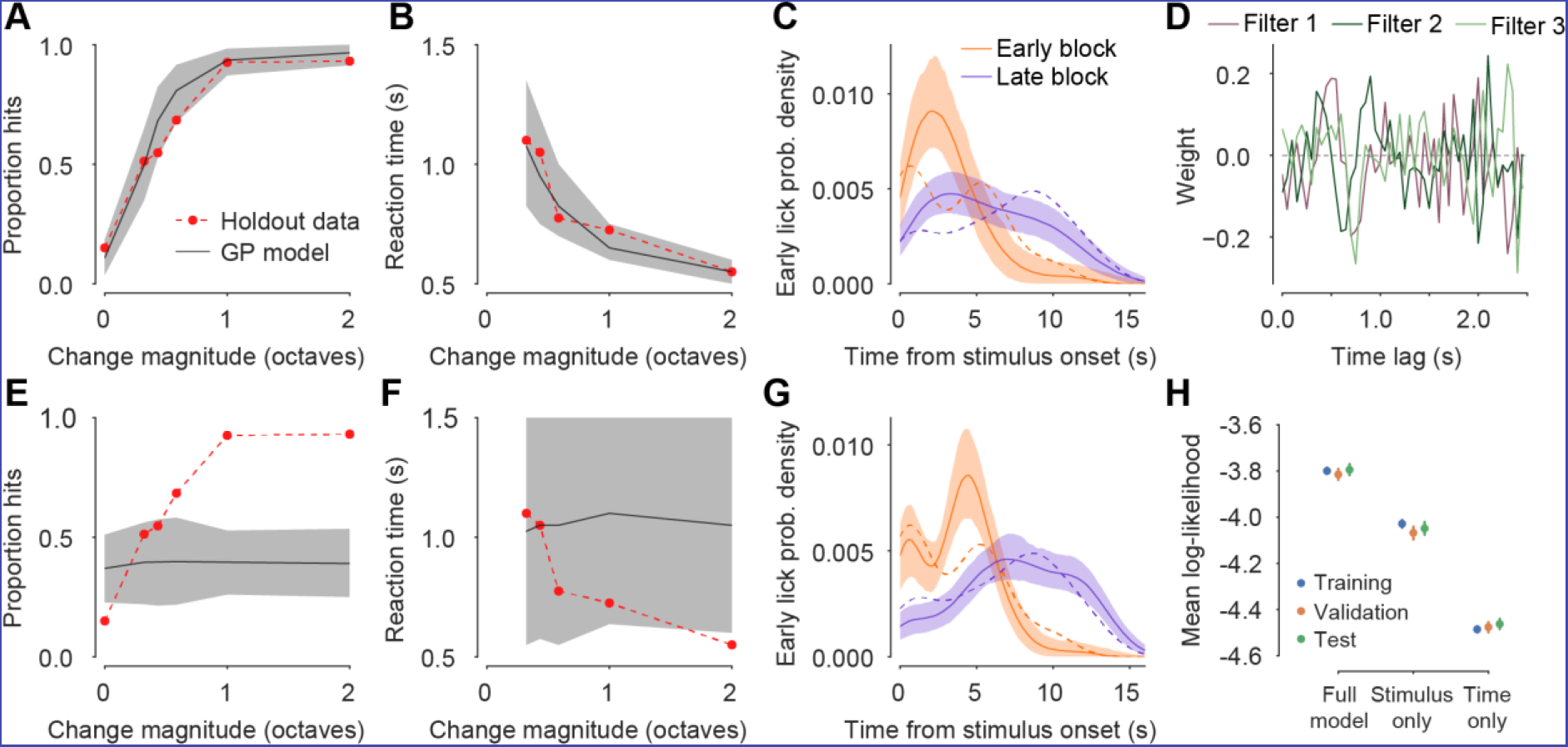
Related to Figure 2. The full stimulus and elapsed time model outperforms models that receive stimulus or time inputs alone. **A-D.** A model that only receives stimulus information captures psychometric (A) and chronometric (B) performance but does not fully account for the timing of early licks (C) or identify both stimulus filters (D). Notation as in Figure 2C-F. **E-G.** A model that only receives timing information does not account for animal’s psychometric (E) and chronometric (F) performance. **H.** The full model outperforms the stimulus only or time only models, as quantified by mean log-likelihood. Error bars – 95% confidence interval, estimated by resampling trials.

**Figure S2.**
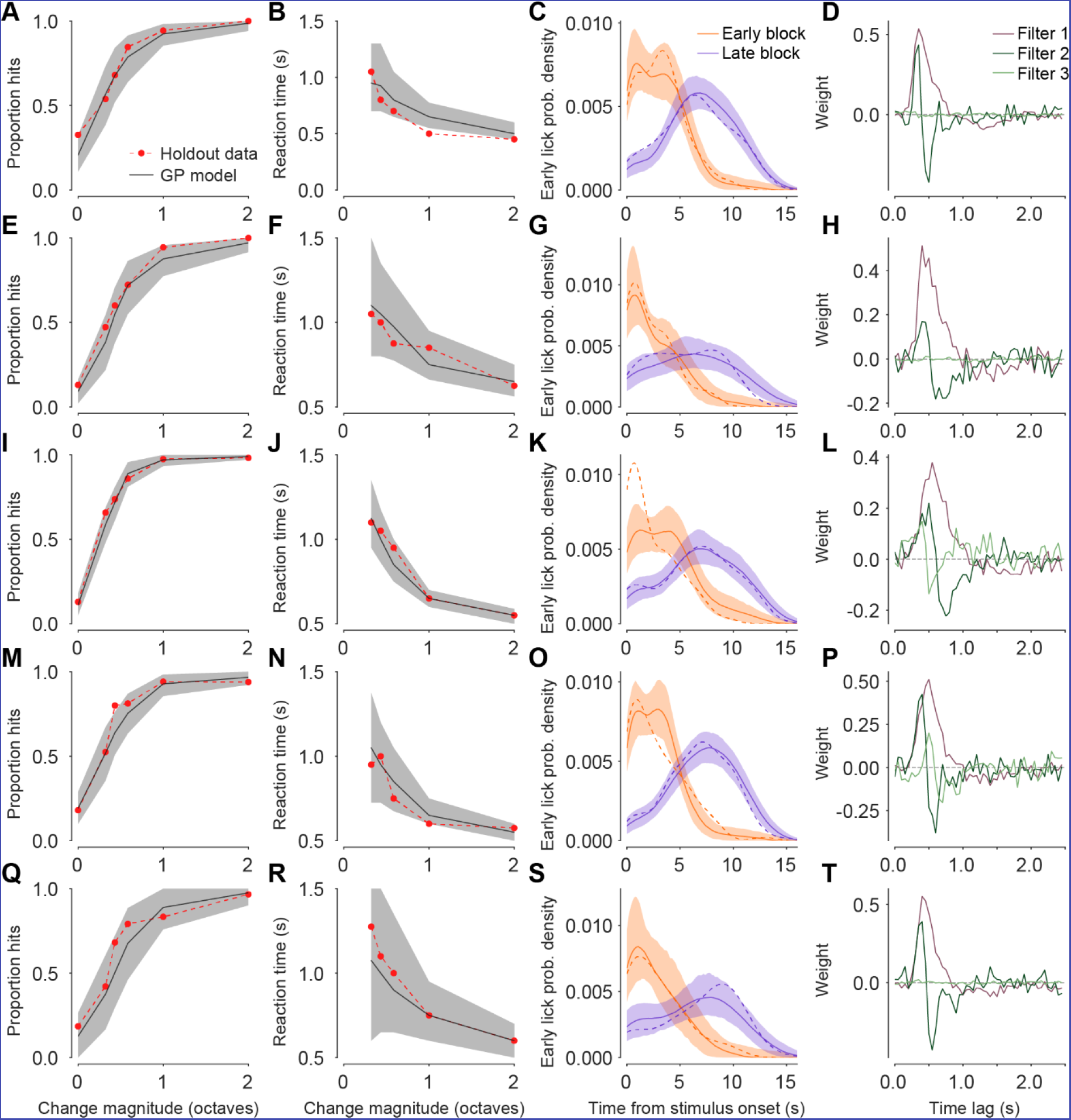
Related to Figure 2. GP classification model captures mouse behavior and identifies similar stimulus filters across mice. Model predictions and holdout data for 5 additional mice; notation as in Figure 2C-F. Columns show psychometric curves (**A, E, I, M, Q**), chronometric curves (**B, F, J, N, R**), early lick timing distributions (**C, G, K, O, S**) and the top three stimulus history filters (**D, H, L, P, T**).

**Figure S3.**
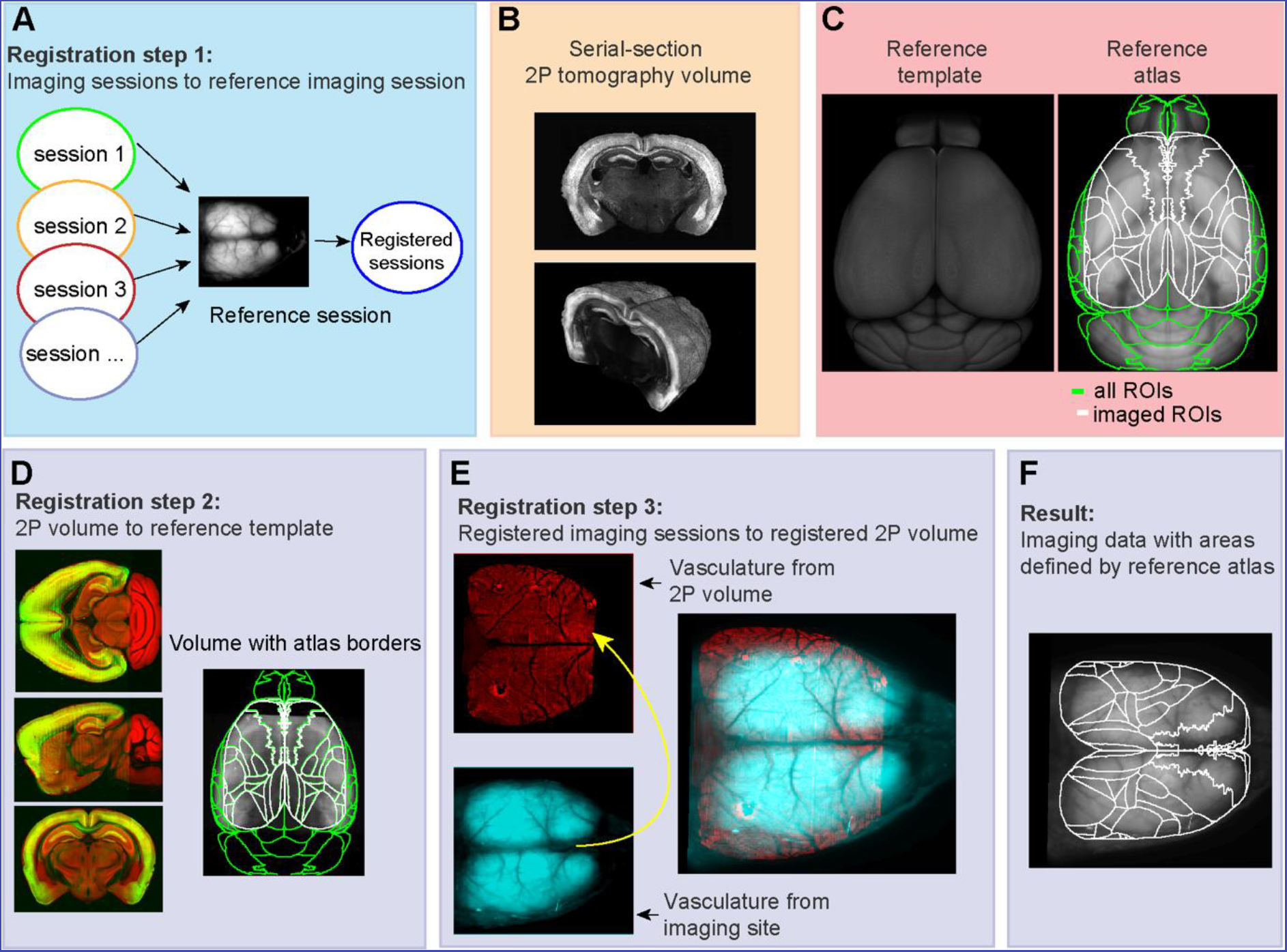
Registration of the imaging data to the reference atlas. **A**. All imaging sessions were registered to a reference imaging session using the vasculature pattern. Each brain has one reference imaging session. **B**. After the *in vivo* imaging experiments, brains were serially sectioned and imaged using a two-photon microscope. Imaged frames were tiled to reconstruct sections (top), and sections were reconstructed to 3D volumes (bottom). C. Reference brain template dorsal view (left) and projection of reference atlas (right) were obtained from the Allen Brain Institute. Borders of all dorsal areas are shown in green. In white are borders trimmed to the extent of the imaging site. D. The sample volume from **B** (in green) was registered to reference template from **C** (in red). Right - atlas projection overlay over projection of registered two-photon volume. E. Vasculature pattern of the reference imaging session (**A**) was registered to the vasculature pattern of the registered two-photon volume from **D**. **F**. The resulting transformation aligned widefield imaging data to the reference atlas coordinate frame.

**Figure S4.**
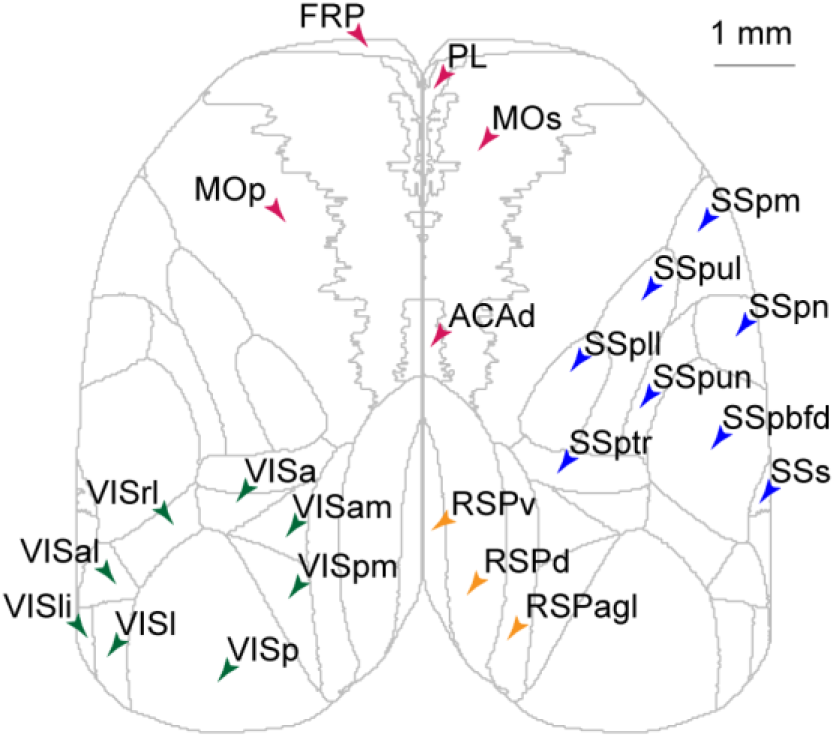
Names and outlines of regions of interest used in this study. VISp: Primary visual area
VISpm: Posteromedial visual area
VISam: Anteromedial visual area
VISa: Anterior area
VISrl: Rostrolateral visual area
VISal: Anterolateral visual area
VISli: Laterointermediate area
VISl: Lateral visual area
RSPagl: Retrosplenial area, lateral agranular part
RSPd: Retrosplenial area, dorsal part
RSPv: Retrosplenial area, ventral part
ACAd: Anterior cingulate area
FRP: Frontal pole, cerebral cortex
PL: Prelimbic area
MOs: Secondary motor area
Mop: Primary motor area
SSs: Supplemental somatosensory area
SSp-bfd: Primary somatosensory area, barrel field
SSp-ll: Primary somatosensory area, lower limb
SSp-m: Primary somatosensory area, mouth
SSp-n: Primary somatosensory area, nose
SSP-tr: Primary somatosensory area, trunk
SSP-ul: Primary somatosensory area, upper limb
SSP-un: Primary somatosensory area, unassigned

**Figure S5.**
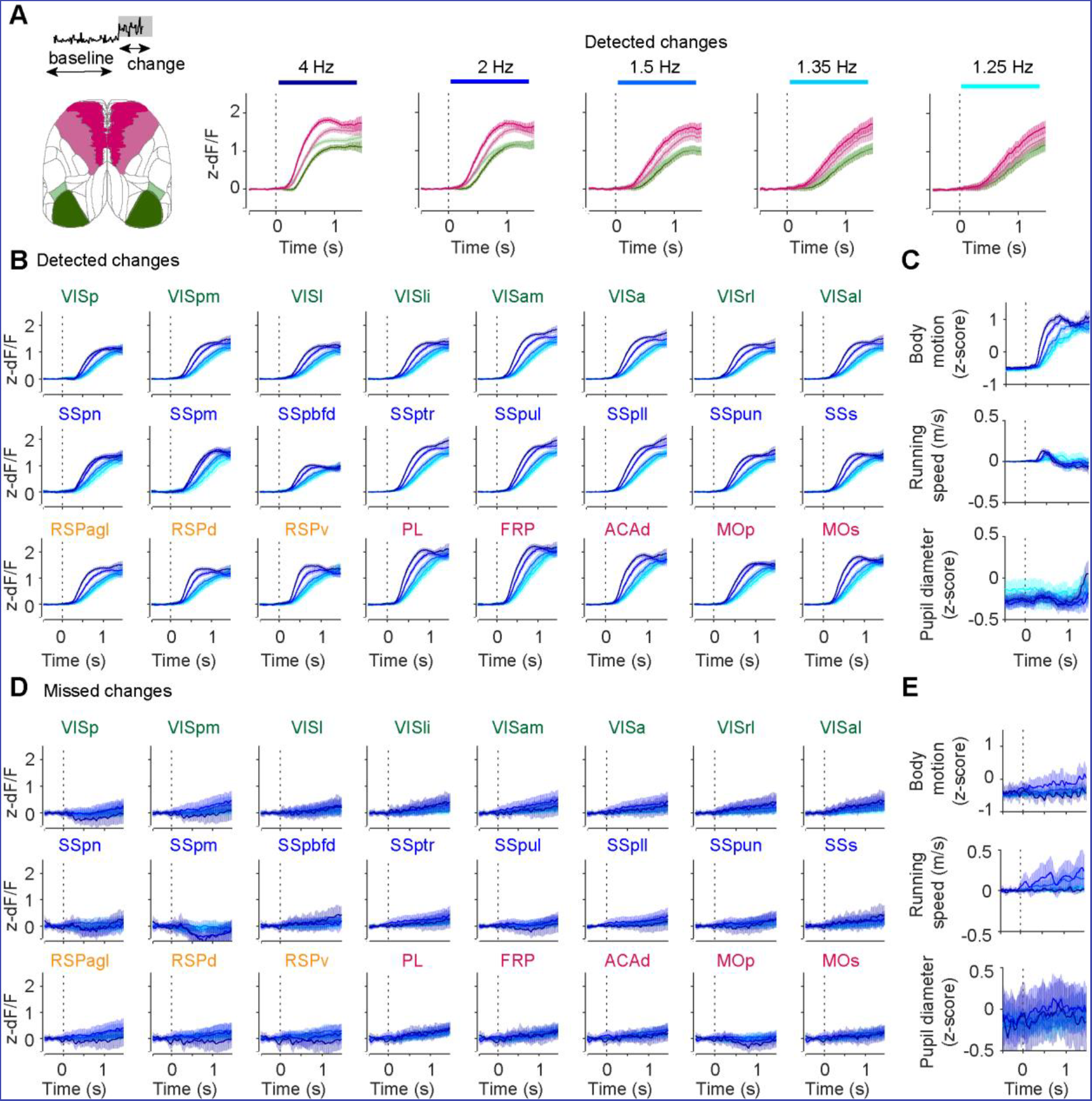
Related to Figure 4D-E. Responses during the change in all imaged regions of interest. **A.** Time courses of responses in VISp, VISrl, MOp, and MOs aligned to change onset on hit trials across the change strengths. **B-C.** Mean z-scored fluorescence of all imaged ROIs (B) and behavioral measures (C) aligned to change onset on hit trials. **D-E.** Mean z-scored fluorescence of all imaged ROIs (D) and behavioral measures (E) aligned to change onset on miss trials.

**Figure S6.**
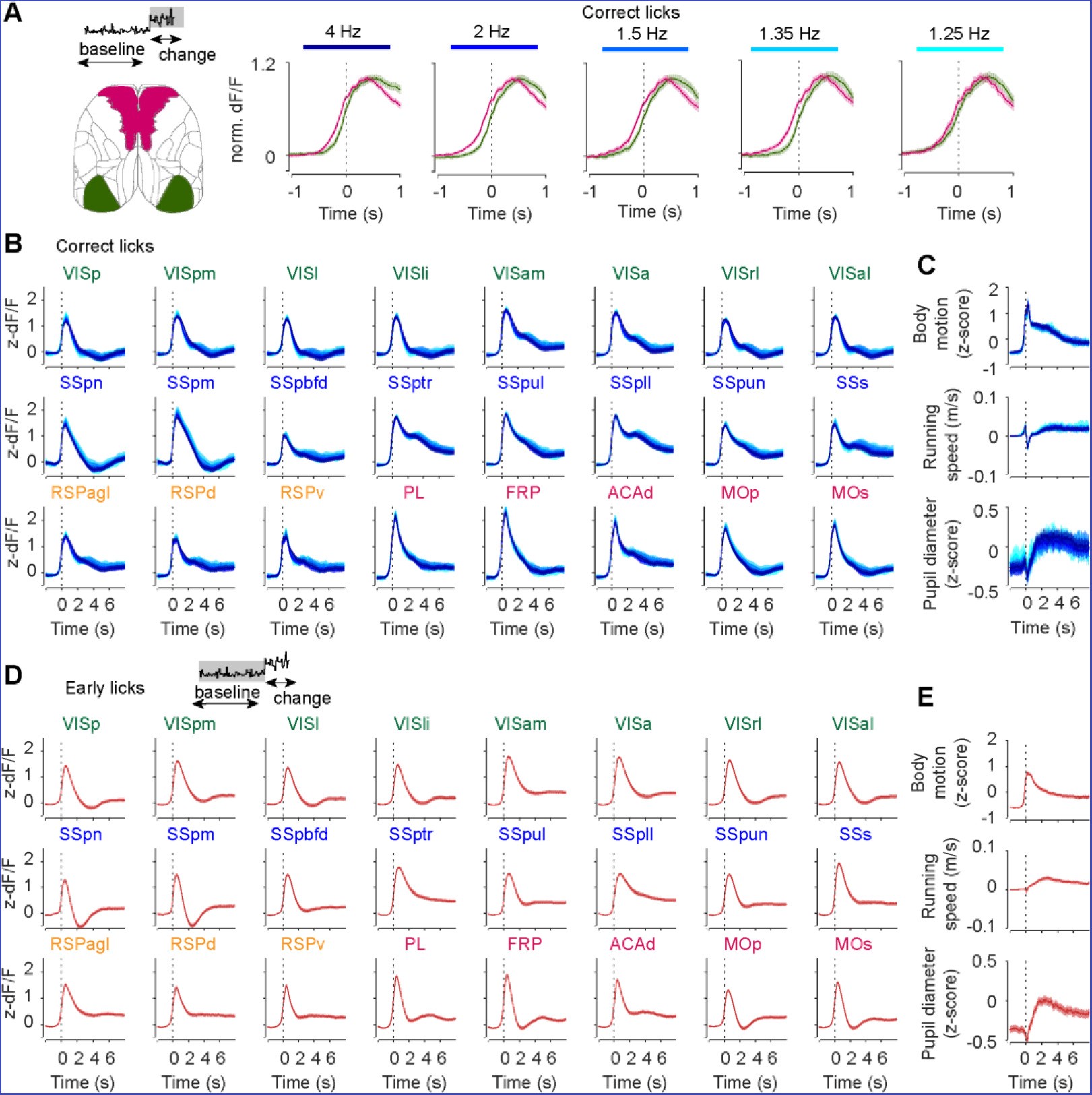
Related to Figure 4E-F. Lick-aligned responses in all imaged regions of interest. **A.** Time courses of responses in VISp and MOs aligned to detection of correct lick across the change strengths. **B-C.** Mean z-scored fluorescence of all imaged ROIs (A) and behavioral measures (B) aligned to licks on hit trials. **D-E.** Mean z-scored fluorescence of all imaged ROIs (C) and behavioral measures (D) aligned to early licks (licks during the baseline period).

**Figure S6.**
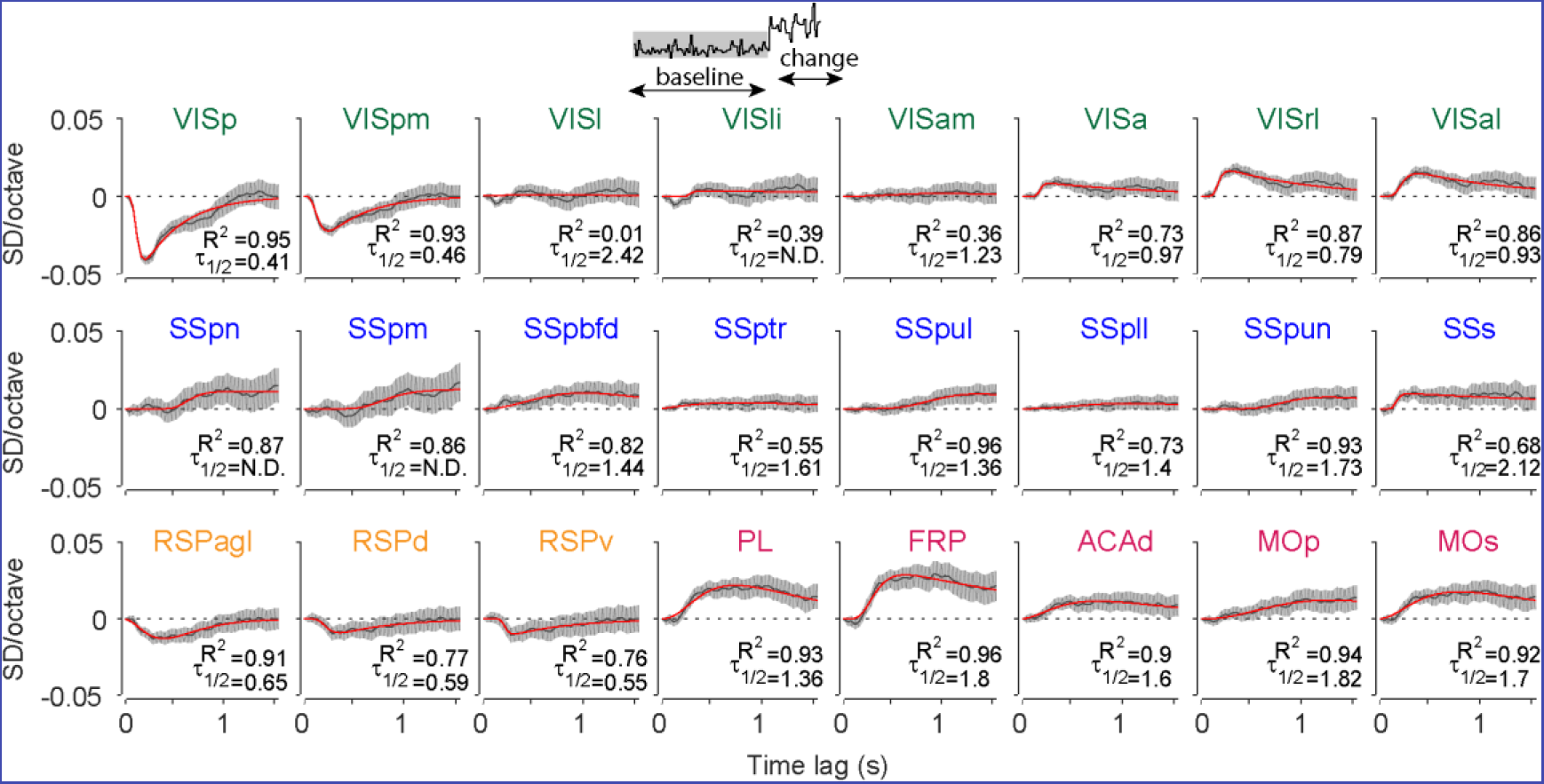
Related to Figure 5B. Responses to subthreshold stimulus fluctuations in all imaged regions of interest. Time course of regression coefficients of widefield fluorescence against baseline stimulus TF in all imaged ROIs; regression coefficients (gray, 95% CI) and multiexponential fits (red).

**Figure S7.**
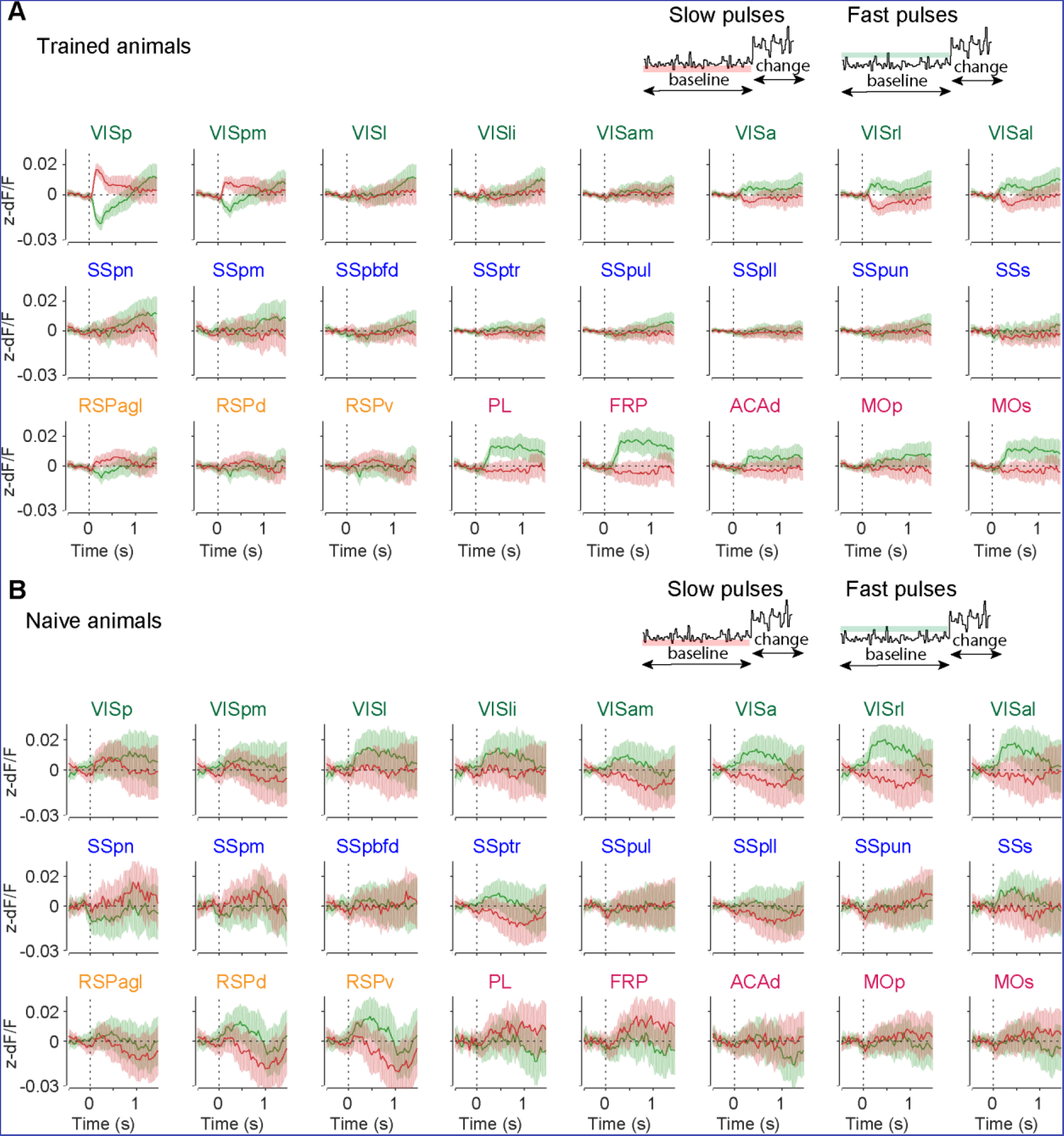
Related to Figure 6B. Responses to anti- and pro-licking subthreshold stimulus fluctuations in all imaged regions of interest in trained and naïve animals. Mean z-scored fluorescence of all imaged ROIs aligned to anti-licking (slow pulses, red) and pro-licking (fast pulses, green) subthreshold stimulus fluctuations in **A.** trained (fast pulses: N = 41194 frames, slow pulses: 42253 frames, 6894 trials, 47 sessions, 6 mice) and **B.** naive mice (fast pulses: N = 14462 frames, slow pulses: 14674 frames, 1680 trials, 10 sessions, 3 mice). Shading is 95% CI.

**Figure S8.**
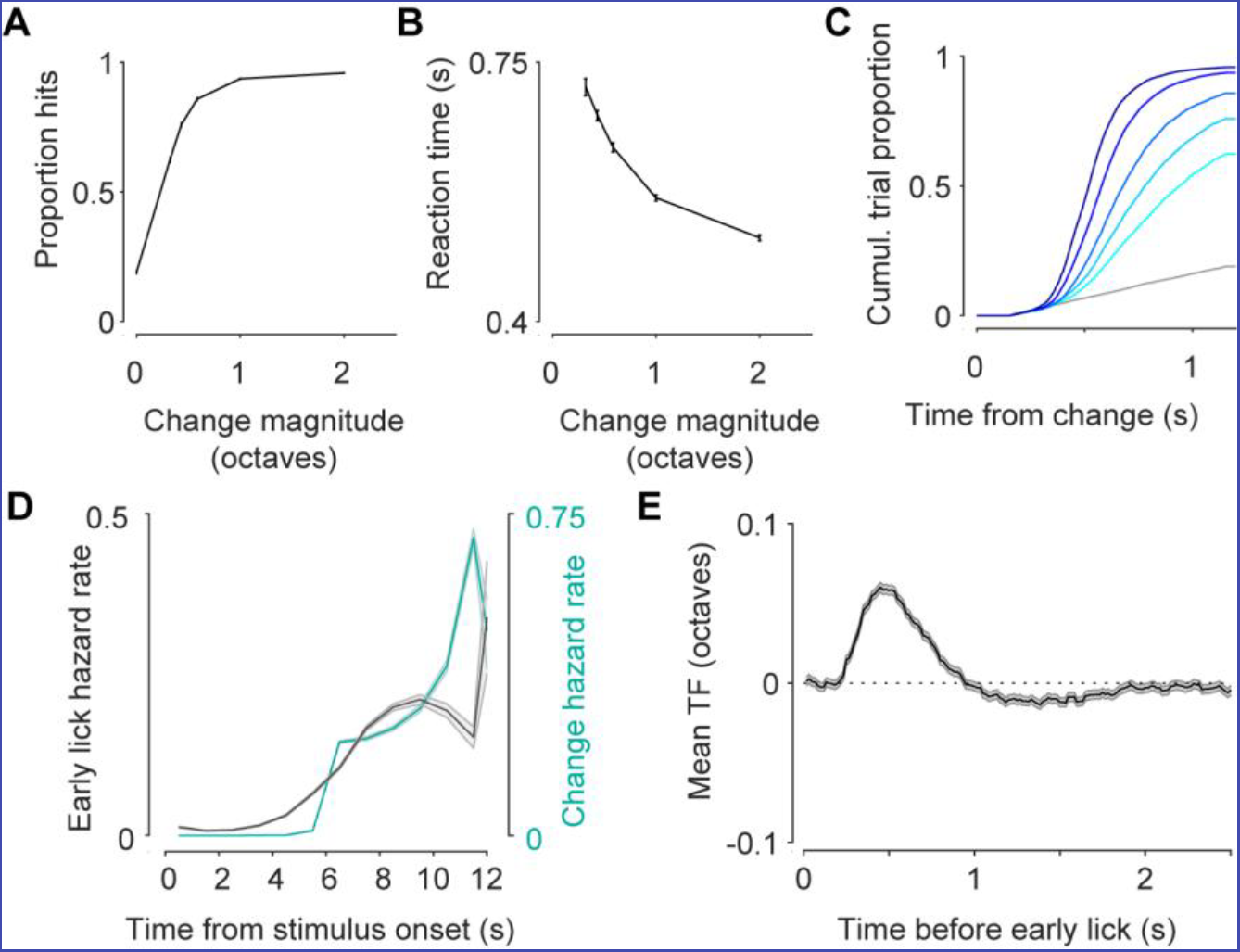
Performance of animals in the running version of the task. **A-B.** Detection rate (**A**) and median reaction times (**B**) are modulated by change magnitude (6 mice, error bars are 95% CI). A. Distribution of reaction times across stimulus changes. Colors same as in Figure 1. B. Early lick hazard rate (gray) and change hazard rate (cyan). E. Average stimulus TF preceding licks during the baseline stimulus (6 mice, shading is 95% CI).

**Figure S9.**
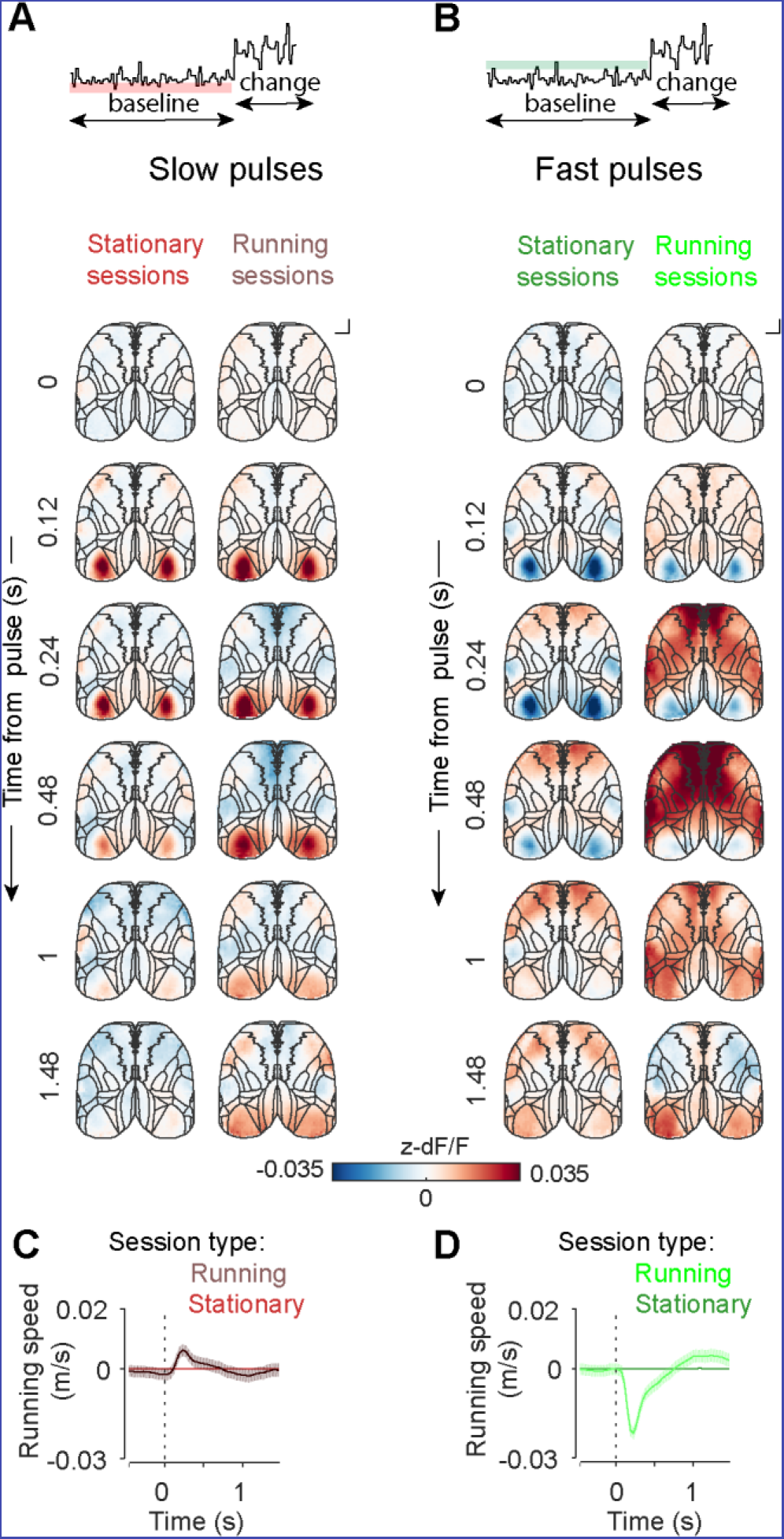
Comparison of responses to fast and slow pulses between running and stationary versions of the task. **A.-B** Maps of mean z-scored fluorescence responses to anti-licking (slow pulses, **A**) and pro-licking (fast pulses, **B**) subthreshold stimulus fluctuations during sessions when mice were required to remain stationary (left). (Fast pulses: N = 41194 frames, slow pulses: 42253 frames, 6894 trials, 47 sessions, 6 mice) or free to run on the wheel (right). (Fast pulses: N = 42004 frames, slow pulses: 42870 frames, reference pulses: 481756, 5930 trials, 37 sessions, 6 mice). C. Slow pulses in running but not stationary mice are followed by an increase in running speed. D. Fast pulses in running but not stationary mice are followed by a reduction in running speed.

**Figure S10.**
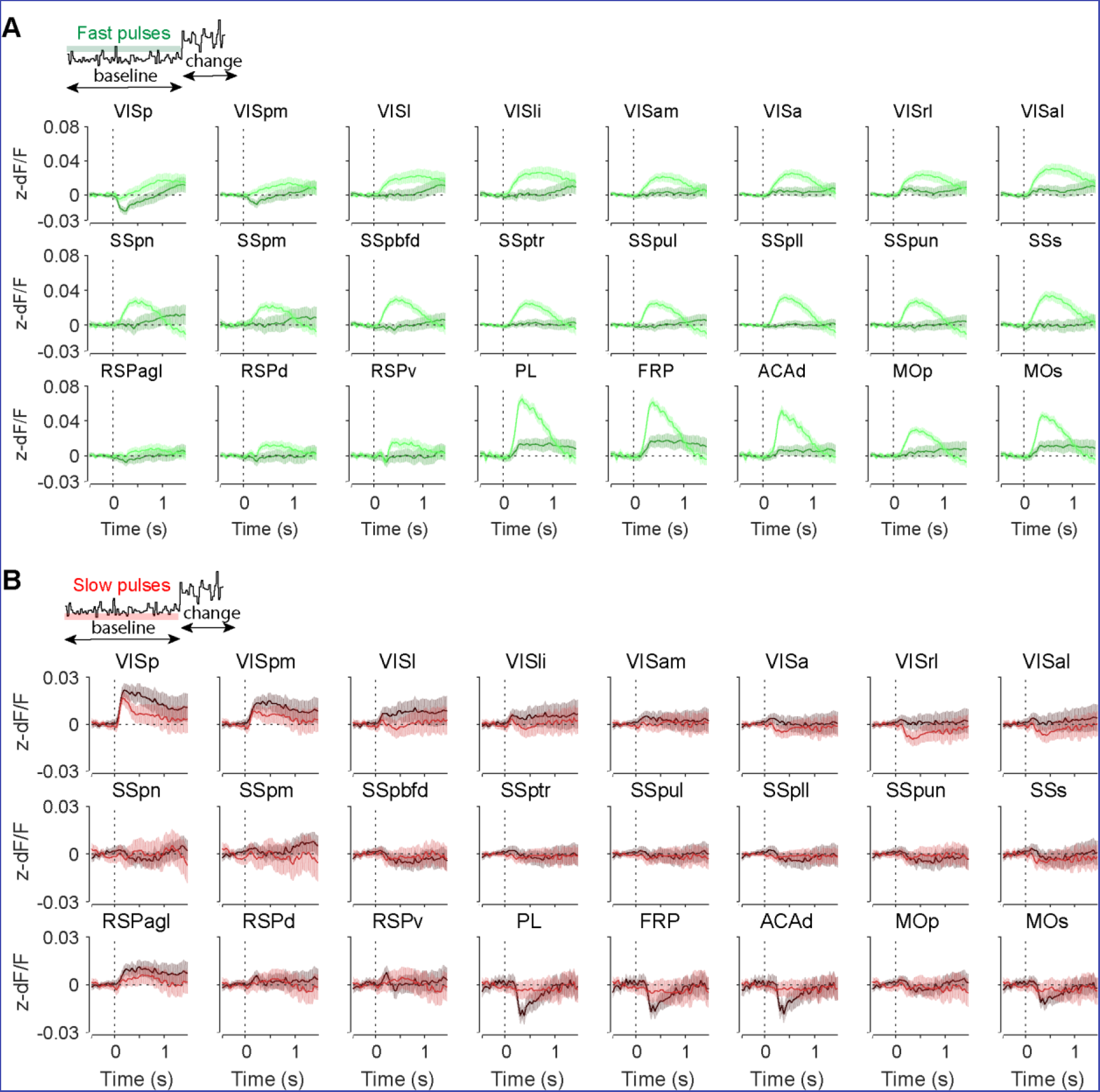
Comparison of responses to fast and slow pulses during running and stationary sessions across all imaged ROIs. **A**. Mean z-scored fluorescence of all imaged ROIs aligned to pro-licking (fast) subthreshold stimulus fluctuations during sessions when mice were required to remain stationary (dark green) or free to run on the wheel (light green). **B.** Mean z-scored fluorescence of all imaged ROIs aligned to anti-licking (slow) subthreshold stimulus fluctuations during sessions when mice were required to remain stationary (red) or free to run on the wheel (black).

**Supplemental Movie 1. Related to Figure 3.** Maps of mean z-scored fluorescence responses aligned to onset of the baseline stimulus in trained (left) and naïve (right) animals. Scalebar 1mm. Playing 0.5x speed.

**Supplemental Movie 2. Related to Figure 4.** Maps of mean z-scored fluorescence responses aligned to change onset on hit trials, sorted by stimulus strength (left to right: 1.25, 1.35, 1.5, 2, and 4 Hz). Scalebar 1mm. Playback at 0.5x speed.

**Supplemental Movie 3. Related to Figure 4.** Maps of mean z-scored fluorescence responses aligned to hit licks, sorted by stimulus strength (left to right: 1.25, 1.35, 1.5, 2, and 4 Hz). Scalebar 1 mm. Playback at 0.5x speed.

**Supplemental Movie 4. Related to Figure 5.** Maps of regression coefficients of widefield fluorescence against baseline stimulus temporal frequency in trained (left) and naïve (right) animals. Scalebar 1 mm. Playback at 0.25x speed.

**Supplemental Movie 5. Related to Figure 6.** Maps of mean z-scored fluorescence responses to pro-(fast, left) and anti-licking (slow, right) subthreshold stimulus fluctuations in trained mice (6 mice). Scalebar 1 mm. Playback at 0.25x speed.

**Supplemental Movie 6. Related to Figure S9B.** Maps of mean z-scored fluorescence responses to pro-(fast) subthreshold stimulus fluctuations in trained mice (6 mice) in stationary (left) and running (right) version of the task. Scalebar 1 mm. Playback at 0.25x speed.

