## Supplemental Information for "Mesoscale cortical dynamics reflect the interaction of sensory evidence and temporal expectation during perceptual decision-making"

### Gaussian process classification model of change detection

#### Model structure

We aim to predict whether and when the mouse would lick in response to the visual stimulus on individual trials of the task. Since the trial is terminated once the mouse licks, we accomplish this by modelling the lick hazard rate – the probability of licking conditioned on the fact that the mouse has not licked up to that point during the trial. If we discretise time during each individual trial into "samples", we will only have the opportunity to observe a lick in the $i$th sample of a trial, if there have been no licks in each of the $i-1$ preceding samples. Therefore, we will represent each trial outcome as the vector $\mathbf{y}$ – a series of zeros terminated by 1 or 0 depending on whether the mouse licked on the particular trial. The likelihood of observing a particular trial outcome $\mathbf{y}$ is the product of conditional likelihoods over the whole trial:

$$p(\mathbf{y}|\mathbf{X}) = p(y_n|y_{n-1}=0,\mathbf{x}_n)\prod_{i=2}^{n-1} p(y_i=0|y_{i-1}=0,\mathbf{x}_i)\,p(y_1=0|\mathbf{x}_1), \quad (1)$$

given the design matrix $\mathbf{X} = [\mathbf{x}_1,\ldots,\mathbf{x}_n]$ containing inputs over the course of the trial. Each input vector $\mathbf{x}_i$ combines the stimulus history in the $Q$ preceding samples and the time elapsed since the stimulus onset:

$$\mathbf{x}_i = [s_i, s_{i-1}, \ldots, s_{i-Q+1}, t_i]^\top. \quad (2)$$

For convenience, we sample the behavior every 50 ms – the duration of individual stimulus fluctuations in the task.

To capture the relationship between the inputs $\mathbf{X}$ and behavior observations $\mathbf{y}$, we assume a latent function $f$ representing the log-odds of licking. Therefore the lick hazard rate at the $i$th moment in time is the logistic function of $f(\mathbf{x}_i)$:

$$p(y_i=1|y_{i-1}=0,\mathbf{x}_i) = p(y_i=1|y_{i-1}=0,f(\mathbf{x}_i)) = \frac{1}{1+\exp(-f(\mathbf{x}_i))}. \quad (3)$$

In order to minimize assumptions on $f$, we do not impose a specific parametrization of $f$ but introduce a prior distribution over functions, using a Gaussian process (GP) with a mean function $m(\mathbf{x})$ and covariance function $\kappa(\mathbf{x},\mathbf{x}')$:

$$f(\mathbf{x}) \sim GP(m(\mathbf{x}), \kappa(\mathbf{x},\mathbf{x}')). \quad (4)$$

The GP prior distribution implies that the log-odds of licking at different moments in time during the session are jointly Gaussian, with the covariance defined as a function the stimulus immediately preceding each moment and time since trial onset:

$$p(f(\mathbf{x}_i), f(\mathbf{x}_j)) = \mathcal{N}\left(\begin{bmatrix} m(\mathbf{x}_i) \\ m(\mathbf{x}_j)\end{bmatrix}, \begin{bmatrix}\kappa(\mathbf{x}_i,\mathbf{x}_i) & \kappa(\mathbf{x}_i,\mathbf{x}_j) \\ \kappa(\mathbf{x}_j,\mathbf{x}_i) & \kappa(\mathbf{x}_j,\mathbf{x}_j)\end{bmatrix}\right) \quad \forall i,j \in [1;n]. \quad (5)$$

In this framework, given a training data set $\mathcal{D} = \{(\mathbf{x}_i, y_i)\}_{i=1}^N$ aggregating all samples for the training trials, we can predict the mouse behavior at any sampled time point $n$ of a test trial using the predictive distribution:

$$p(\mathbf{y}_*|\mathbf{X}_*,\mathcal{D}) = \int p(y_{*n}|y_{*n-1}=0,f_{*n})\prod_{i=2}^{n-1}p(y_{*i}=0|y_{*i-1}=0,f_{*i})\,p(y_{*1}=0|f_{*1})\,p(\mathbf{f}_*|\mathcal{D})\,d\mathbf{f}_* \tag{6}$$

where $\mathbf{X}_* = [\mathbf{x}_{*1},\ldots,\mathbf{x}_{*n}]$ is the test input data, $\mathbf{f}_* = [f_{*1},\ldots,f_{*n}]^\top = [f(x_{*1}),\ldots,f(x_{*n})]^\top$ the vector of corresponding latent function values and $\mathbf{y}_* = [y_{*1},\ldots,y_{*n}]^\top$ the behavior whose probability is evaluated.

The next section will describe how to evaluate the posterior distribution of $f$ and what it implies in terms of training. The approach is based on the work of Hensman et al. (2015) and is summarized here for the sake of completeness and reproducibility. The subsequent sections will describe the kernel and mean functions and summarize specifics of the implementation.

#### GP posterior estimation

The posterior distribution can be rewritten as an integral of two terms:

$$p(\mathbf{f}_*|\mathcal{D}) = \int p(\mathbf{f}_*|\mathbf{f})\,p(\mathbf{f}|\mathcal{D})\,d\mathbf{f}, \tag{7}$$

where $\mathbf{f} = [f_1,\ldots,f_N]$ is the vector of latent function values on the training data. The first term is the conditional distribution of the latent function values for the test data given the latent function values of the training data. As a property of the GP prior, we can analytically derive its form as the density of a multivariate normal distribution, that we note $p(\mathbf{f}_*|\mathbf{f}) = \mathcal{N}(\boldsymbol{\mu}_*,\boldsymbol{\Sigma}_*)$. The mean and covariance parameters $(\boldsymbol{\mu}_*,\boldsymbol{\Sigma}_*)$ depend on the covariance of the training data $[\mathbf{K}_{\mathbf{ff}}]_{i,j} = \kappa(\mathbf{x}_i,\mathbf{x}_j)$, and the covariance of the test data with the training data $[\mathbf{K}_{*\mathbf{f}}]_{i,j} = \kappa(\mathbf{x}_{*i},\mathbf{x}_j)$:

$$\boldsymbol{\mu}_* = m(\mathbf{X}_*) + \mathbf{K}_{*\mathbf{f}}\mathbf{K}_{\mathbf{ff}}^{-1}(\mathbf{f}-m(\mathbf{X})) \quad \text{and} \quad \boldsymbol{\Sigma}_* = \mathbf{K}_{\mathbf{ff}} - \mathbf{K}_{*\mathbf{f}}\mathbf{K}_{\mathbf{ff}}^{-1}\mathbf{K}_{*\mathbf{f}}^\top, \tag{8}$$

where $m(\mathbf{X}_*) = [m(\mathbf{x}_{*1}),\ldots,m(\mathbf{x}_{*n})]^\top$ and $m(\mathbf{X}) = [m(\mathbf{x}_1),\ldots,m(\mathbf{x}_N)]^\top$.

The second term $p(\mathbf{f}|\mathcal{D})$, the posterior of the latent function values on the training data, is more problematic. As our likelihood is not normally distributed (Eq. 3), it cannot be expressed in closed-form (Rasmussen 2006). We replace it with a normal variational approximating distribution $q(\mathbf{f}) = \mathcal{N}(\boldsymbol{\mu}_\mathbf{f},\boldsymbol{\Sigma}_\mathbf{f})$. The mean and covariance parameters $(\boldsymbol{\mu}_\mathbf{f},\boldsymbol{\Sigma}_\mathbf{f})$ are optimized to minize the Kullback-Leibler (KL) divergence between $p(\mathbf{f}|\mathcal{D})$ and $q(\mathbf{f})$, a measure of discrepancy between the two distributions.

With this variational approximation, we can replace Eq. 7 with an approximate posterior distribution:

$$p(\mathbf{f}_*|\mathcal{D}) \approx q(\mathbf{f}_*|\mathcal{D}) = \int p(\mathbf{f}_*|\mathbf{f})\,q(\mathbf{f})\,d\mathbf{f}, \tag{9}$$

which possesses a closed-form expression. Both terms in the integral being normal distribution density functions, the result is also a normal distribution density $q(\mathbf{f}_*|\mathcal{D}) = \mathcal{N}(\tilde{\boldsymbol{\mu}}_*,\tilde{\boldsymbol{\Sigma}}_*)$ with parameters defined as follows:

$$\tilde{\boldsymbol{\mu}}_* = m(\mathbf{X}_*) + \mathbf{K}_{*\mathbf{f}}\mathbf{K}_{\mathbf{ff}}^{-1}(\boldsymbol{\mu}_\mathbf{f}-m(\mathbf{X})) \quad \text{and} \quad \tilde{\boldsymbol{\Sigma}}_* = \mathbf{K}_{\mathbf{ff}} - \mathbf{K}_{*\mathbf{f}}(\mathbf{K}_{\mathbf{ff}}-\boldsymbol{\Sigma}_\mathbf{f})^{-1}\mathbf{K}_{*\mathbf{f}}^\top. \tag{10}$$

For a given test input $\mathbf{x}_{*i}$, the posterior mean can be rewritten as:

$$\tilde{\mu}_{*i} = m(\mathbf{x}_{*i}) + \sum_{j=1}^{N}\alpha_i\kappa(\mathbf{x}_*,\mathbf{x}_j) \quad \text{where} \quad \boldsymbol{\alpha} = \mathbf{K}_{\mathbf{ff}}^{-1}(\boldsymbol{\mu}_\mathbf{f}-m(\mathbf{X})). \tag{11}$$

Making inferences using Eq. 10 requires the entire training data set $\mathcal{D}$ and becomes computationally intractable for large $N$. To reduce computational complexity, we replace the training data with a set of $M$ (such that $M \ll N$) pseudo input points, also called inducing points, $\mathbf{Z} = [\mathbf{z}_1, \ldots, \mathbf{z}_N]$ and latent function values $\mathbf{u} = [f(\mathbf{z}_1), \ldots, f(\mathbf{z}_M)]$. In this scenario, the variational approximating distribution becomes $q(\mathbf{f}) = \int p(\mathbf{f}|\mathbf{u})q(\mathbf{u})\, d\mathbf{u}$ where $q(\mathbf{u}) = \mathcal{N}(\boldsymbol{\mu}_\mathbf{u}, \boldsymbol{\Sigma}_\mathbf{u})$. With this new variational approximation, the parameters of the approximate posterior distribution in Eq. 10 become:

$$\tilde{\boldsymbol{\mu}} = m(\mathbf{X}_*) + \mathbf{K}_{*\mathbf{u}}\mathbf{K}_{\mathbf{uu}}^{-1}(\boldsymbol{\mu}_\mathbf{u} - m(\mathbf{Z})) \quad \text{and} \quad \tilde{\boldsymbol{\Sigma}} = \mathbf{K}_{\mathbf{uu}} - \mathbf{K}_{*\mathbf{u}}(\mathbf{K}_{\mathbf{uu}} - \boldsymbol{\Sigma}_\mathbf{u})^{-1}\mathbf{K}_{*\mathbf{f}}^\top, \tag{12}$$

where $[\mathbf{K}_{\mathbf{uu}}]_{i,j} = \kappa(\mathbf{z}_i, \mathbf{z}_j)$ and $[\mathbf{K}_{*\mathbf{u}}]_{i,j} = \kappa(\mathbf{x}_{*i}, \mathbf{z}_j)$. As a consequence, any computation involving these quantities only requires inverting a $M \times M$ matrix, rather than a $N \times N$ matrix.

In this final formulation, training a model consists in optimizing the values of the inducing points and values $(\mathbf{Z}, \mathbf{u})$ as well as the variational distribution parameters $(\boldsymbol{\mu}_\mathbf{u}, \boldsymbol{\Sigma}_\mathbf{u})$, in order to minimize the KL-divergence between $p(\mathbf{f}|\mathcal{D})$ and $q(\mathbf{f})$. Minimizing the KL-divergence objective is equivalent to maximizing the model evidence lower bound (ELBO) defined as:

$$\begin{aligned} \mathcal{L} &= \mathbb{E}_{q(\mathbf{f})}[\log p(\mathbf{y}|\mathbf{f})] - \mathrm{KL}(\mathrm{q}(\mathbf{u})||\mathrm{p}(\mathbf{u})) \\ &= \sum_{i=1}^{N} \mathbb{E}_{q(\mathbf{f})}[\log p(y_i|y_{i-1}, f_i)] - \mathrm{KL}(\mathrm{q}(\mathbf{u})||\mathrm{p}(\mathbf{u})). \end{aligned} \tag{13}$$

As the ELBO approximates the model marginal log-likelihood, it is also used to learn the model hyperparameters, i.e. the kernel function and mean function parameters. All parameters are optimized using a gradient descent technique. The definition of the ELBO involving a simple sum over the training data, we employ a stochastic version the gradient descent using only a random subset of the training data at each iteration.

#### Kernel and mean functions

In general the mean function $m(\mathbf{x})$ can be set to 0 with no reduction in model performance, as the posterior distribution can capture the mean. However, as the mean log-odds of licking are far below 0 (mice do not lick for the vast majority model samples), we include a constant mean function to ensure that the model makes sensible predictions outside of the range of the training data.

We define the kernel function as the sum of stimulus and time dependent components:

$$\kappa(\mathbf{x}, \mathbf{x}') = \kappa_s(\mathbf{x}, \mathbf{x}') + \kappa_t(\mathbf{x}, \mathbf{x}'), \tag{14}$$

where $\kappa_s(\mathbf{x}, \mathbf{x}')$ and $\kappa_t(\mathbf{x}, \mathbf{x}')$ depend on stimulus history $\mathbf{s}$ or the time elapsed since the stimulus onset $t$, respectively. An advantage of the additive form of the kernel is that stimulus- and time-dependent components of the log-odds can be readily separated. The posterior predictive mean from Eq. 11 can be decomposed into:

$$\begin{aligned} \tilde{\mu}_{*i} &= m(\mathbf{x}_{*i}) + \sum_{j=1}^{N} \alpha_i(\kappa_s(\mathbf{x}_{*i}, \mathbf{x}_j) + \kappa_t(\mathbf{x}_{*i}, \mathbf{x}_j)) \\ &= m(\mathbf{x}_{*i}) + \sum_{j=1}^{N} \alpha_i\kappa_s(\mathbf{x}_{*i}, \mathbf{x}_j) + \sum_{j=1}^{N} \alpha_i\kappa_t(\mathbf{x}_{*i}, \mathbf{x}_j) \\ &= m(\mathbf{x}_{*i}) + \tilde{\mu}_{is} + \tilde{\mu}_{it}\ . \end{aligned} \tag{15}$$

To identify stimulus features that best explain observed behavior, we first multiply the stimulus history by a $Q \times D$ matrix of filters $\mathbf{W}$ (Snelson 2006, Vivarelli & Williams 1999), where $Q$ is the number of stimulus history samples included in the model and $D$ is the number of stimulus filters:

$$\boldsymbol{\phi} = \mathbf{s}^\top \mathbf{W}. \tag{16}$$

Our approach is to select an arbitrarily large $D$ and control the effective dimensionality of the filtered stimulus space by placing an automatic relevance determination (ARD) prior on $\mathbf{W}$ (Beal 2003, Bishop 1999). The ARD prior assumes a zero-mean Gaussian prior on each column $\mathbf{w}_d$ of the matrix $\mathbf{W}$:

$$p(\mathbf{W}|\boldsymbol{\nu}) = \prod_{d=1}^{D} \left(\frac{\nu_d}{2\pi}\right)^{D/2} \exp\left(-\frac{\nu_d ||\mathbf{w}_d||^2}{2}\right), \tag{17}$$

and a gamma distributed prior on the precision vector $\boldsymbol{\nu} = [\nu_1, \ldots, \nu_D]$. This prior over $\boldsymbol{\nu}$ favors high precisions, consequently shrinking the columns of $\mathbf{W}$ that do not contribute to the prediction performance of the model. To estimate the loadings of $\mathbf{W}$, we infer an approximate posterior distribution over $(\mathbf{W}, \boldsymbol{\nu})$ using the automatic differentiation variational inference (ADVI) framework (Kucukelbir et al. 2017), which extends the ELBO definition of the model (Eq. 13) with additional terms.

We then use a Matérn 5/2 kernel over filter outputs $\boldsymbol{\phi}$ such that $\mathbf{x} = [\phi_1, \ldots, \phi_D, t]^\top$. Since their magnitude can be adjusted by scaling the columns of $\mathbf{W}$, the length scale of the stimulus kernel is fixed to 1 to avoid over-parametrizing the model. Therefore, $\kappa_s$ depends only on the euclidean distance $L$ between $\boldsymbol{\phi}$ and $\boldsymbol{\phi}'$:

$$\kappa_s(\mathbf{x}, \mathbf{x}') = \sigma_s^2 (1 + \sqrt{5}L + 5L/3) \exp(-\sqrt{5}L), \text{ where } L = ||\boldsymbol{\phi} - \boldsymbol{\phi}'||_2. \tag{18}$$

It is well established that precision of timing behavior is not constant but varies with the duration of time intervals. To account for this non-stationarity, we passed the input to the time component of the kernel through a non-linear monotonic warping function, parametrized as a sum of $\tanh$ functions (Snelson et al. 2004):

$$t_w = t + \sum_{k=1}^{J} a_k \tanh(b_k(t + c_k)). \tag{19}$$

We optimize the parameters $(\mathbf{a}, \mathbf{b}, \mathbf{c})$ during model training. The time kernel $\kappa_t$ is then computed as a Matérn 5/2 kernel over warped time with its own length scale and variance parameters:

$$\kappa_t(\mathbf{x}, \mathbf{x}') = \sigma_t^2 \left(1 + \frac{\sqrt{5}(t_w - t'_w)}{\ell_t} + \frac{5(t_w - t'_w)^2}{3\ell_t^2}\right) \exp\left(-\frac{\sqrt{5}(t_w - t'_w)}{\ell_t}\right). \tag{20}$$

#### Hierarchical structure

Our data set contains trials recorded under different experimental conditions, such as blocks of trials with different distributions of stimulus change times, as well as running and stationary animals. We aimed to extend the model to capture the differences in behavior between these experimental blocks, while also learning their shared features. The GP framework offers a simple and rigorous approach for dealing with such structured data (Hensman et al. 2013).

We introduce an indicator variable $b$, which denotes the experimental block for each sample, and split each part of the covariance $\kappa(\mathbf{x}, \mathbf{x}')$ into population and block-specific components:

$$\kappa(\mathbf{x}, \mathbf{x}') = \begin{cases} \kappa_{s_p} + \kappa_{s_b} + \kappa_{t_p} + \kappa_{t_b} & \text{when } b = b' \\ \kappa_{s_p} + \kappa_{t_p} & \text{otherwise.} \end{cases} \tag{21}$$

Population and block-specific covariance functions share the same forms, described by Eqs. 18-20, and stimulus features defined by $\mathbf{W}$ but have their own hyperparameters, variances $\sigma_s^2$ and $\sigma_t^2$, and length scale $\ell_t$.

### Model implementation and training

The model is implemented on the basis of the Stochastic Variational GP class of the *GPflow* Python package (de G. Matthews et al. 2017), which relies on *Tensorflow* (Abadi et al. 2015) for automatic differentiation and GPU-based computations.

Model training was carried out independently for each mouse, using behavioral data from both stationary and trained versions of the task. The block variable $b$ (Eq. 21) indicated whether a trial was acquired in the running task, or in early or late hazard rate blocks in the stationary task. The data set was split into training (60%), validation (20%), and test (20%) sets, stratified according to change strength and experimental block.

We used 450 inducing points and assigned 150 to each experimental block. Inducing points were initialized using a mini-batch variant of the K-means clustering algorithm, provided by the *scikit-learn* Python package (Buitinck et al. 2013). We set the number of stimulus history samples $Q$ to 50, the number of filters $D$ to 15, and the number of $\tanh$ functions $J$ (Eq. 19) to 5. Filter coefficients $\mathbf{W}$ and the parameters $(\mathbf{a}, \mathbf{b}, \mathbf{c})$ were randomly initialized from a standard normal distribution.

We performed the optimization of all parameters using the Adam algorithm (Kingma & Ba 2014) with the default settings and a learning rate of 0.001 employing 12000 samples per mini-batch. Computations took place on Nvidia GeForce 2070 RTX or 2080 RTX graphics cards, and were terminated after a maximum time of 10 hours or when convergence was reached as assessed by predictive performance on the validation set.

Once a model was fitted, we estimated the predictive distribution for each time point of each trial with Eq. 6, replacing the integral with a Monte Carlo integral using 500 samples from the approximate posterior distribution. We did not use the full posterior distribution of $W$ but replaced it with the posterior mean. To assess the performance of the model and capture uncertainty of its predictions, we generated behavior replicates by drawing licks from the predictive distribution for each trial in the test set. If a sampled lick occurred after the baseline period, the corresponding replicate trial was labeled as a hit and a reaction time was calculated as the time from change to the lick. We repeated this sampling procedure to obtain 500 replicates of the test data set and generated psychometric and chronometric curves and the distribution of lick times for each replicate. Figure 2 and Figures S1-2 show the median, 2.5%, and 97.5% quantiles of the sampled curves. We also performed model comparisons based on their predictive performance, using the averaged predictive log-likelihood of trials in the data set (Figure S1H).
